# Microbial function and genital inflammation in young South African women at high risk of HIV infection

**DOI:** 10.1101/2020.03.10.986646

**Authors:** Arghavan Alisoltani, Monalisa T. Manhanzva, Matthys Potgieter, Christina Balle, Liam Bell, Elizabeth Ross, Arash Iranzadeh, Michelle du Plessis, Nina Radzey, Zac McDonald, Bridget Calder, Imane Allali, Nicola Mulder, Smritee Dabee, Shaun Barnabas, Hoyam Gamieldien, Adam Godzik, Jonathan M. Blackburn, David L. Tabb, Linda-Gail Bekker, Heather B. Jaspan, Jo-Ann S. Passmore, Lindi Masson

**Affiliations:** Division of Medical Virology, Department of Pathology, University of Cape Town, Cape Town 7925, South Africa; Division of Biomedical Sciences, University of California Riverside School of Medicine, Riverside, CA 92521, USA; Computational Biology Division, Department of Integrative Biomedical Sciences, University of Cape Town, Cape Town 7925, South Africa; Division of Chemical and Systems Biology, Department of Integrative Biomedical Sciences, University of Cape Town, Cape Town 7925, South Africa; Division of Immunology, Department of Pathology, University of Cape Town, Cape Town 7925, South Africa; Centre for Proteomic and Genomic Research, Cape Town 7925, South Africa; Laboratory of Human Pathologies Biology, Department of Biology and Genomic Center of Human Pathologies, Mohammed V University in Rabat, Morocco; Institute of Infectious Disease and Molecular Medicine (IDM), University of Cape Town, Cape Town 7925, South Africa; Centre for Infectious Diseases Research (CIDRI) in Africa Wellcome Trust Centre, University of Cape Town, Cape Town 7925, South Africa; Seattle Children’s Research Institute, University of Washington, WA 98101, USA; Bioinformatics Unit, South African Tuberculosis Bioinformatics Initiative, Stellenbosch University, Stellenbosch 7602, South Africa; DST–NRF Centre of Excellence for Biomedical Tuberculosis Research, Stellenbosch University, Stellenbosch 7602, South Africa; Desmond Tutu HIV Centre, Cape Town 7925, University of Cape Town, South Africa; Centre for the AIDS Programme of Research in South Africa, Durban 4013, South Africa; National Health Laboratory Service, Cape Town 7925, South Africa; Disease Elimination Program, Life Sciences Discipline, Burnet Institute, Melbourne 3004, Australia; Central Clinical School, Monash University, Melbourne 3004, Australia

**Keywords:** Metaproteomics, microbiome, microbial function, female genital tract, inflammation, cytokine

## Abstract

**Background:** Female genital tract (FGT) inflammation is an important risk factor for HIV acquisition. The FGT microbiome is closely associated with inflammatory profile, however, the relative importance of microbial activities has not been established. Since proteins are key elements representing actual microbial functions, this study utilized metaproteomics to evaluate the relationship between FGT microbial function and inflammation in 113 young and adolescent South African women at high risk of HIV infection. Women were grouped as having low, medium or high FGT inflammation by K-means clustering according to pro-inflammatory cytokine concentrations.

**Results:** A total of 3,186 microbial and human proteins were identified in lateral vaginal wall swabs using liquid chromatography-tandem mass spectrometry, while 94 microbial taxa were included in the taxonomic analysis. Both metaproteomics and 16S rRNA gene sequencing analyses showed increased non-optimal bacteria and decreased lactobacilli in women with FGT inflammatory profiles. However, differences in the predicted relative abundance of most bacteria were observed between 16S rRNA gene sequencing and metaproteomics analyses. Bacterial protein functional annotations (gene ontology) predicted inflammatory cytokine profiles more accurately than bacterial relative abundance determined by 16S rRNA gene sequence analysis, as well as functional predictions based on 16S rRNA gene sequence data (p<0.0001). The majority of microbial biological processes were underrepresented in women with high inflammation compared to those with low inflammation, including a *Lactobacillus*-associated signature of reduced cell wall organization and peptidoglycan biosynthesis. This signature remained associated with high FGT inflammation in a subset of 74 women nine weeks later, was upheld after adjusting for *Lactobacillus* relative abundance, and was associated with *in vitro* inflammatory cytokine responses to *Lactobacillus* isolates from the same women. Reduced cell wall organization and peptidoglycan biosynthesis were also associated with high FGT inflammation in an independent sample of ten women.

**Conclusions:** Both the presence of specific microbial taxa in the FGT and their properties and activities are critical determinants of FGT inflammation. Our findings support those of previous studies suggesting that peptidoglycan is directly immunosuppressive, and identify a possible avenue for biotherapeutic development to reduce inflammation in the FGT. To facilitate further investigations of microbial activities, we have developed the FGT-METAP application that is available at (http://immunodb.org/FGTMetap/).

## Background

Despite the large reduction in AIDS-related deaths as a result of the expansion of HIV antiretroviral programs, global HIV incidence has declined by only 16% since 2010 [1]. Of particular concern are the extremely high rates of HIV infection in young South African women [1]. The behavioral and biological factors underlying this increased risk are not fully understood, however, a key predisposing factor that has been identified in this population is genital inflammation [2–4]. We have previously shown that women with elevated concentrations of pro-inflammatory cytokines and chemokines in their genital tracts were at higher risk of becoming infected with HIV [2]. Furthermore, the protection offered by a topical antiretroviral tenofovir microbicide was compromised in women with evidence of genital inflammation [3].

The mechanisms by which female genital tract (FGT) inflammation increases HIV risk are likely multifactorial, and increased genital inflammatory cytokine concentrations may facilitate the establishment of a productive HIV infection by recruiting and activating HIV target cells, directly promoting HIV transcription and reducing epithelial barrier integrity [5–7]. Utilizing proteomics, Arnold, *et al*. showed that proteins associated with tissue remodeling processes were up-regulated in the FGT, while protein biomarkers of mucosal barrier integrity were down-regulated in individuals with an inflammatory profile [7]. This suggests that inflammation increases tissue remodeling at the expense of mucosal barrier integrity, which may in turn increase HIV acquisition risk [7]. Furthermore, neutrophil-associated proteins, especially certain proteases, were positively associated with inflammation and may be involved in disrupting the mucosal barrier [7]. This theory was further validated by the finding that antiproteases associated with HIV resistance were increased in a cohort of sex workers from Kenya [8].

Bacterial vaginosis (BV), non-optimal cervicovaginal bacteria, and sexually transmitted infections (STIs), which are highly prevalent in South African women, are likely important drivers of genital inflammation in this population [9,10]. BV and non-optimal bacteria have been consistently associated with a marked increase in pro-inflammatory cytokine concentrations [9–12]. Conversely, women who have “optimal” vaginal microbiota, primarily consisting of *Lactobacillus* spp., have low levels of genital inflammation and a reduced risk of acquiring HIV [13]. Lactobacilli and the lactic acid that they produce may actively suppress inflammatory responses and thus may play a significant role in modulating immune profiles in the FGT [14–16]. However, partly due to the complexity and diversity of the microbiome, the immunomodulatory mechanisms of specific vaginal bacterial species are not fully understood [17]. Further adding to this complexity is the fact that substantial differences in the cervicovaginal microbiota exist by geographical location and ethnicity and that the properties of different strains within particular microbial species are also highly variable [15,18].

In addition to providing insight into host mucosal barrier function, metaproteomic studies of the FGT can evaluate microbial activities and functions that may influence inflammatory profiles [19]. A recent study highlighted the importance of vaginal microbial function by demonstrating that FGT bacteria, such as *Gardnerella vaginalis*, modulate the efficacy of the tenofovir microbicide by active metabolism of the drug [20]. Additionally, Zevin et al. found that bacterial functional profiles were associated with epithelial barrier integrity and wound healing in the FGT [21], suggesting that microbial function may also be closely linked to genital inflammation. The aim of the present study was to utilize both metaproteomics and 16S rRNA gene sequencing to improve our understanding of the relationship between microbial function and inflammatory profiles in the FGTs of South African women.

## Results

This study included 113 young and adolescent HIV-uninfected women (aged 16-22 years) residing in Cape Town, South Africa [22]. Seventy-four of these women had samples and metadata available for analysis at a second time-point nine weeks later. Ten women had samples available at the second time-point, but not the first, and were included to validate the functional signature identified in the primary analysis of this study. STI and BV prevalence were high in this cohort, with 48/113 (42%) of the women having at least one STI and 56/111 (50%) of the women being BV positive by Nugent scoring (Additional file 1: Table S1). The use of injectable contraceptives, BV, and chlamydia were associated with increased genital inflammatory cytokines, as previously described [10,23].

### FGT metaproteome associates with BV and inflammatory cytokine profiles

Liquid chromatography-tandem mass spectrometry (LC-MS/MS) analysis was conducted on the total protein acquired from lateral vaginal wall swabs for all participants. Using principal component analysis (PCA), it was found that women with BV clustered separately from women who were BV negative according to the relative abundance of all host and microbial proteins identified (Additional file 1: Fig. S1a). Women who were considered to have high levels of genital inflammation based on pro-inflammatory cytokine concentrations (Additional file 1: Fig. S2) tended to cluster separately from women with low inflammation, while no clear grouping was observed by STI status or chemokine profiles (Additional file 1: Fig. S1b-d). After adjusting for potentially confounding variables including age, contraceptives, prostate specific antigen (PSA) and co-infections, BV was associated with the largest number of differentially abundant proteins, followed by inflammatory cytokine profiles, while fewer associations were observed between chemokine profiles and protein relative abundance and none between STI status and protein relative abundance (Additional file 1: Fig. S1e). Individual STIs were also not associated with marked changes in the metaproteome, with *Chlamydia trachomatis, Neisseria gonorrhoeae, Trichomonas vaginalis* and *Mycoplasma genitalium* associated with significant changes in zero, five, seven and eleven proteins, respectively (after adjustment for potential confounders and multiple comparisons). However, this stratified analysis is limited by the small number of STI cases for some infections and the prevalence of co-infections.

### Differences in taxonomic assignment using metaproteomics and 16S rRNA gene sequencing

Of the proteins identified in lateral vaginal wall swab samples, 38.8% were human, 55.8% were bacterial, 2.7% fungal, 0.4% archaeal, 0.09% of viral origin, and 2% grouped as “other” (not shown). However, as the majority of taxa identified had <2 proteins detected, a more stringent cut-off was applied to include only taxa with >3 detected proteins, or 2 proteins detected in multiple taxa. The final curated dataset included 44% human, 55% bacterial (n=81 taxa), and 1% fungal proteins (n=13 taxa) (Fig. 1a-d; Additional file 2: Table S2). When the relative abundance of the most abundant bacterial taxa identified using metaproteomics was compared to the relative abundance of the most abundant bacteria identified using 16S rRNA gene sequencing, a large degree of similarity was found at the genus level. A total of 6/9 of the genera identified using 16S rRNA gene sequence analysis were also identified using metaproteomics (Fig. 1e, f; Additional file 1: Fig. S3). As expected, both approaches showed that multiple *Lactobacillus* species were less abundant in women with high versus low inflammation, while BV-associated bacteria (including *G. vaginalis, Prevotella* species, *Megasphaera* species, *Sneathia amnii* and *Atopobium vaginae)* were more abundant in women with high inflammation compared to those with low inflammation (Additional file 2: Table S3). However, species-level annotation differed between the 16S rRNA gene sequencing and metaproteomics analyses, with metaproteomics identifying more lactobacilli at the species level, while *Lachnovaginosum* genomospecies (previously known as BVAB1) was not identified using metaproteomics. Additionally, the relative abundance of the taxa differed between 16S rRNA gene sequencing and metaproteomics analyses.

**Figure 1.**
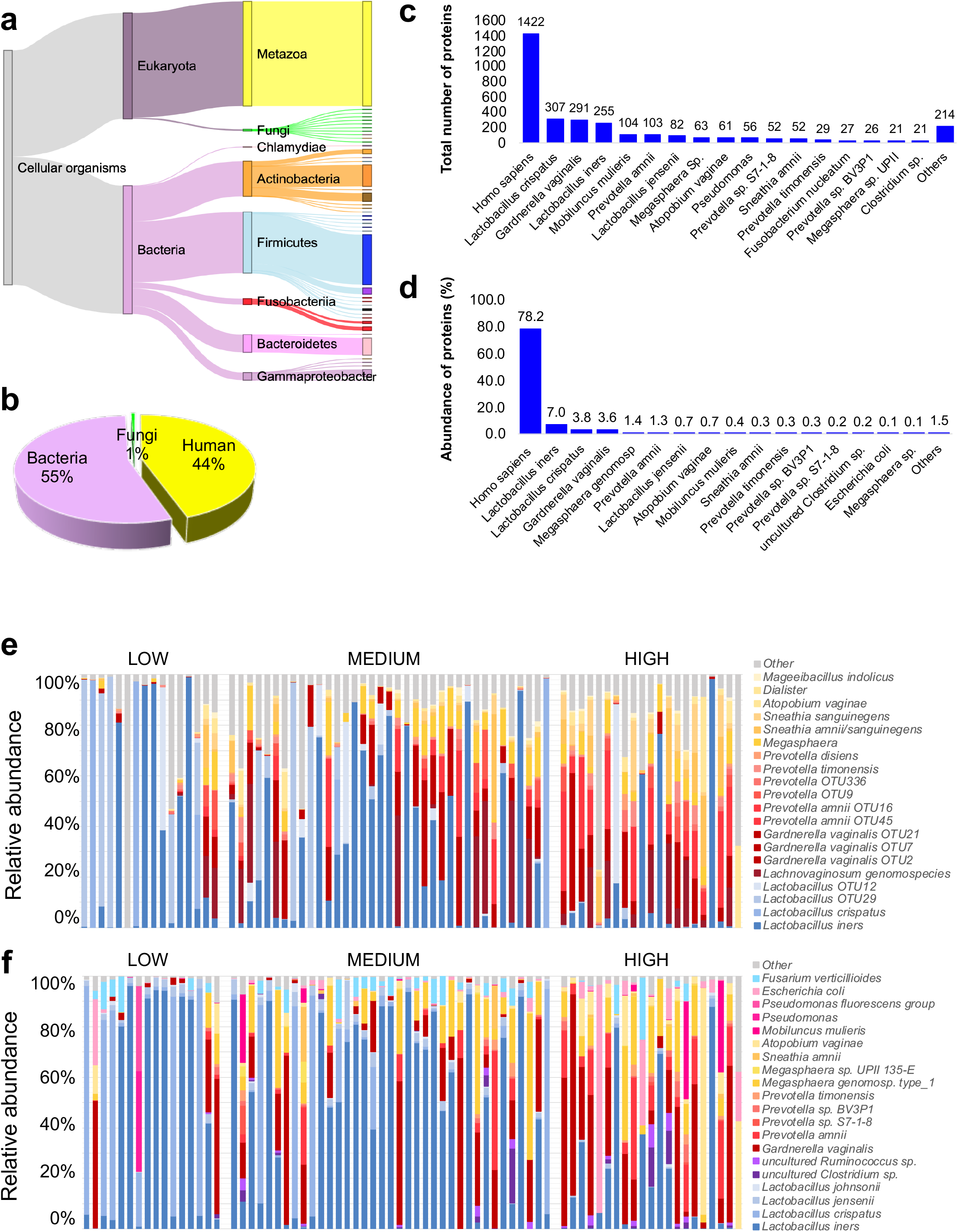
Bacterial relative abundance determined using metaproteomics versus 16S rRNA gene sequencing by inflammation cytokine profile. Liquid chromatography-tandem mass spectrometry was used to evaluate the metaproteome in lateral vaginal wall swabs from 113 women from Cape Town, South Africa. Proteins were identified using MaxQuant and a custom database generated using *de novo* sequencing to filter the UniProt database. Taxonomy was assigned using UniProt and relative abundance of each taxon was determined by aggregating the intensity-based absolute quantification (iBAQ) values of all proteins identified for each taxon. (**a**) Proteins identified were assigned to the Eukaryota and Bacteria domains. Eukaryota proteins included those assigned to the fungi kingdom and metazoan subkingdom, while the bacteria domain included actinobacteria, firmicutes, fusobacteria, bacteriodetes and gammaproteobacter phyla. (**b**) Distribution of taxa identified is shown as a pie chart. (**c**) Number of proteins detected for taxa for which the greatest number of proteins were identified. (**d**) Protein relative abundance per taxon for taxa with the highest relative abundance. The relative abundance of the top 20 most abundant bacterial taxa identified using (**e**) 16S rRNA gene sequencing and (**f**) metaproteomics is shown for all participants for whom both 16S rRNA gene sequence data and metaproteomics data were generated (n=74). For 16S rRNA gene sequence analysis, the V4 region was amplified and libraries sequenced on an Illumina MiSeq platform. Inflammation groups were defined based on hierarchical followed by K-means clustering of all women according to the concentrations of nine pro-inflammatory cytokine concentrations [interleukin (IL)-1α, IL-1β, IL-6, IL-12p40, IL-12p70, tumor necrosis factor (TNF)-α, TNF-β, TNF-related apoptosis-inducing ligand (TRAIL), interferon (IFN)-γ]. OTU: Operational taxonomic unit.

### Protein profiles differ between women defined as having high, medium or low FGT inflammation

A total of 449, 165, and 39 host and microbial proteins were differentially abundant between women with high versus low inflammation, medium versus high inflammation and medium versus low inflammation, respectively (Additional file 2: Table S4). Microbial proteins that were more abundant in women with high or medium inflammation versus low inflammation were mostly assigned to non-optimal bacterial taxa (including *G. vaginalis, Prevotella* species, *Megasphaera* species, *S. amnii* and *A. vaginae*). Less abundant microbial proteins in women with high inflammation were primarily of *Lactobacillus* origin (Additional file 1: Fig. S4). Of the human proteins that were differentially abundant according to inflammation status, 93 were less abundant and 132 were more abundant in women with high compared to those with low inflammation. Human biological process gene ontologies (GOs) that were significantly underrepresented in women with high versus low inflammation included positive regulation of apoptotic signaling, establishment of endothelial intestinal barrier, ectoderm development and cornification [false discovery rate (FDR) adj. p<0.0001 for all; Additional file 1: Fig. S5]. Significantly overrepresented pathways in women with high versus low inflammation included multiple inflammatory processes - such as chronic response to antigenic stimulus, positive regulation of IL-6 production and inflammatory response (FDR adj. p<0.0001 for all; Additional file 1: Fig. S5).

Weighted correlation network analysis of microbial and host proteins identified five modules (clusters) representing co-correlations between microbial and host proteins (Fig. 2a, b). The yellow module included primarily *L. iners* proteins, while the turquoise module primarily consisted of *L. crispatus* and host proteins. The grey and brown modules consisted entirely of host proteins and the blue module included non-optimal bacteria and host proteins. Pro-inflammatory cytokines correlated inversely with the *Lactobacillus* modules (yellow and turquoise) and positively with the grey, brown and blue modules. Chemokines were significantly positively correlated with the brown and grey modules and inversely with the turquoise module (Fig. 2c). Blue, yellow, turquoise and grey modules correlated with BV Nugent score, while BV showed no significant correlation with the brown module (Fig. 2d). The finding that the brown module did not include any microbial proteins and was not associated with BV or STIs, suggests the presence of inflammatory processes independent of the microbiota. For STIs, no significant correlations (p-value <0.05) with any of the five modules were detected (Fig. 2d). To determine whether the co-correlations of microbial proteins with host proteins had functional meaning, we profiled the top biological processes of proteins as shown in the row sidebar in Fig. 2a. Host proteins that correlated negatively with the blue module (consisting of non-optimal bacteria) were mostly involved in cell adhesion, cell-cell adhesion, cell-cell junction assembly, and regulation of cell-adhesion pathways. This suggests reduced epithelial barrier function associated with *G. vaginalis, Prevotella* spp., *Megasphaera*, and *A. vaginae*. Host proteins involved in immune system and inflammatory response pathways were positively correlated with brown and grey modules and inversely associated with the turquoise module including *L. crispatus*. Despite variability in the microbial proteins and taxa, redundant microbial functional pathways were identified across different modules, indicating that microbiota share a set of common metabolic functions (e.g. glucose metabolism, protein translation and synthesis), as expected. Although most lactobacilli proteins were negatively associated with BV and pro-inflammatory cytokines (see turquoise and yellow modules in Fig. 2a), almost all *L. iners* proteins were clustered in the yellow module with slightly different functional pathways from lactobacilli proteins in the turquoise module.

**Figure 2.**
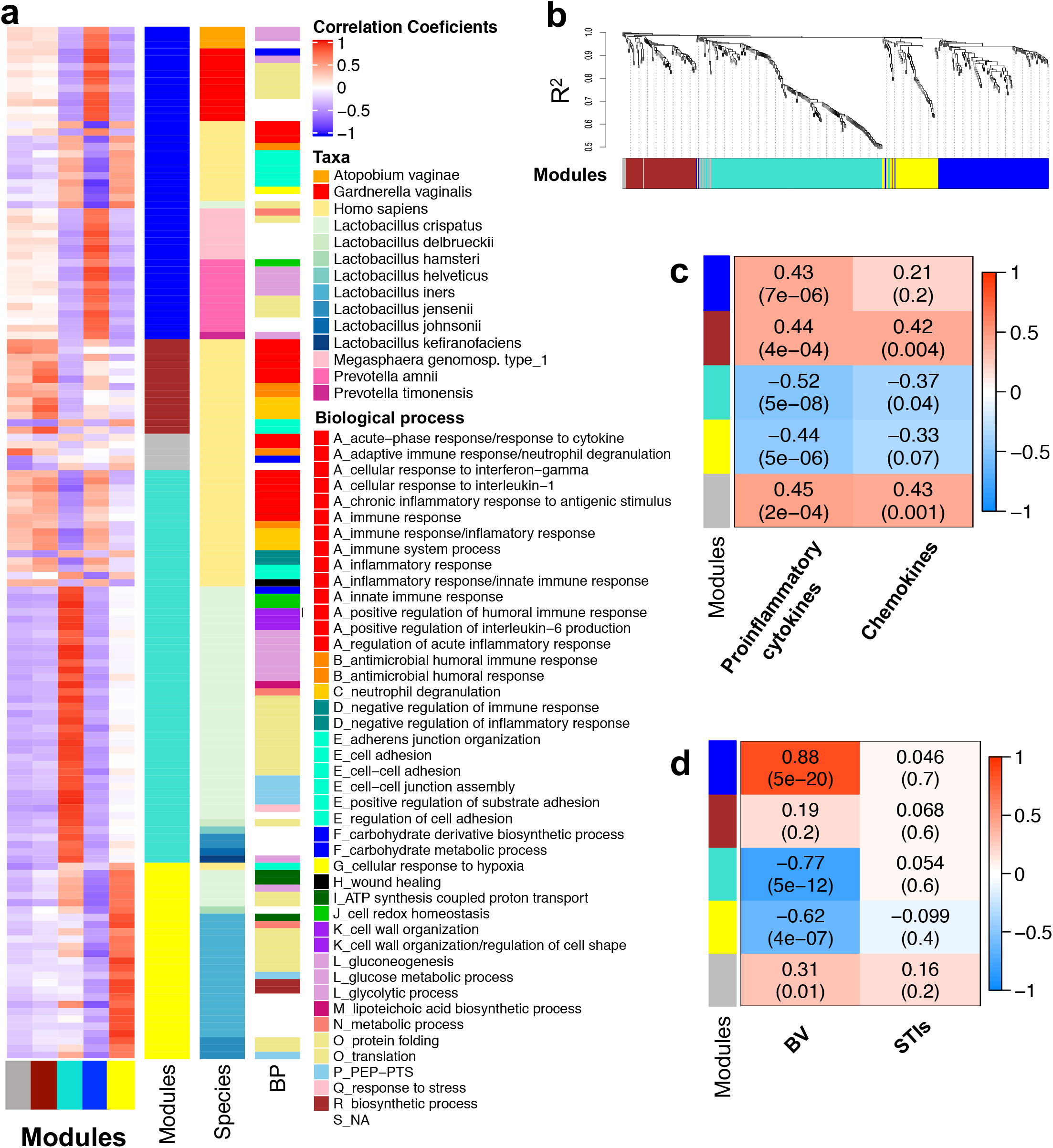
Weighted co-correlation network analysis of microbial, and human proteins. The weighted correlation network analysis (WGCNA) R package was used to build a microbial-host functional weighted co-correlation network using intensity-based absolute quantification (iBAQ) values. **a**) Correlations between proteins and modules are shown by the heatmap, with positive correlations shown in red and negative correlations shown in blue. Row sidebars represent the top taxa and biological processes assigned to each of the proteins (no separate correlation coefficients were calculated for taxa and biological processes). **b**) The protein dendrogram and module assignment are displayed, with five modules identified using dynamic tree cut. **c**) Spearman’s rank correlation was used to determine correlation coefficients between individual cytokines/chemokines and module eigengenes (the first principal component of the expression matrix of the corresponding module) for all samples. Finally, we reported the average Spearman’s correlation coefficients for all pro-inflammatory cytokines and chemokines. **d**) Similarly, Spearman’s correlation coefficients were calculated between module eigengenes and Nugent scores [bacterial vaginosis (BV)] and sexually transmitted infections (STIs) as a categorical variable.

### Microbial function predicts genital inflammation with greater accuracy than taxa relative abundance

The accuracies of (i) microbial relative abundance (determined using 16S rRNA gene sequencing), (ii) functional prediction based on 16S rRNA gene sequence data, (iii) microbial protein relative abundance (determined using metaproteomics) and (iv) microbial molecular function (determined by aggregation of GOs of proteins identified using metaproteomics) for prediction of genital inflammation status (low, medium and high groups) were evaluated using random forest analysis. The three most important species predicting genital inflammation grouping by 16S rRNA gene sequence data were *Sneathia sanguinegens, A. vaginae*, and *Lactobacillus iners* (Fig. 3a). The top functional pathways based on 16S rRNA gene sequence data for the identification of women with inflammation were prolactin signaling, biofilm formation, and tyrosine metabolism (Fig. 3b). Similarly, the most important proteins predicting inflammation grouping by metaproteomics included two *L. iners* proteins (chaperone protein DnaK and pyrophosphate phospho-hydrolase), two *A. vaginae* proteins (glyceraldehyde-3-phosphate dehydrogenase and phosphoglycerate kinase), *Megasphaera* genomosp. type 1 cold-shock DNA-binding domain protein, and *L. crispatus* ATP synthase subunit alpha (Fig. 3c). The top molecular functions associated with genital inflammation included oxidoreductase activity (of multiple proteins expressed primarily by lactobacilli, but also *Prevotella* and *Atopobium* species), beta-phosphoglucomutase activity (expressed by lactobacilli) and GTPase activity (of multiple proteins produced by mainly *Prevotella, G. vaginalis*, and some lactobacilli) as illustrated in Fig. 3d. These findings suggest that the functions of key BV-associated bacteria and lactobacilli are critical for determining the level of genital inflammation. It was also found that the out-of-bag (OOB) error rate distribution was significantly lower for molecular function, followed by protein relative abundance, functional prediction based on 16S rRNA gene sequence data and then taxa relative abundance (Fig. 3e). This suggests that the activities of the bacteria present in the FGT play a critical role in driving genital inflammation, in addition to the presence and relative abundance of particular taxa.

**Figure 3.**
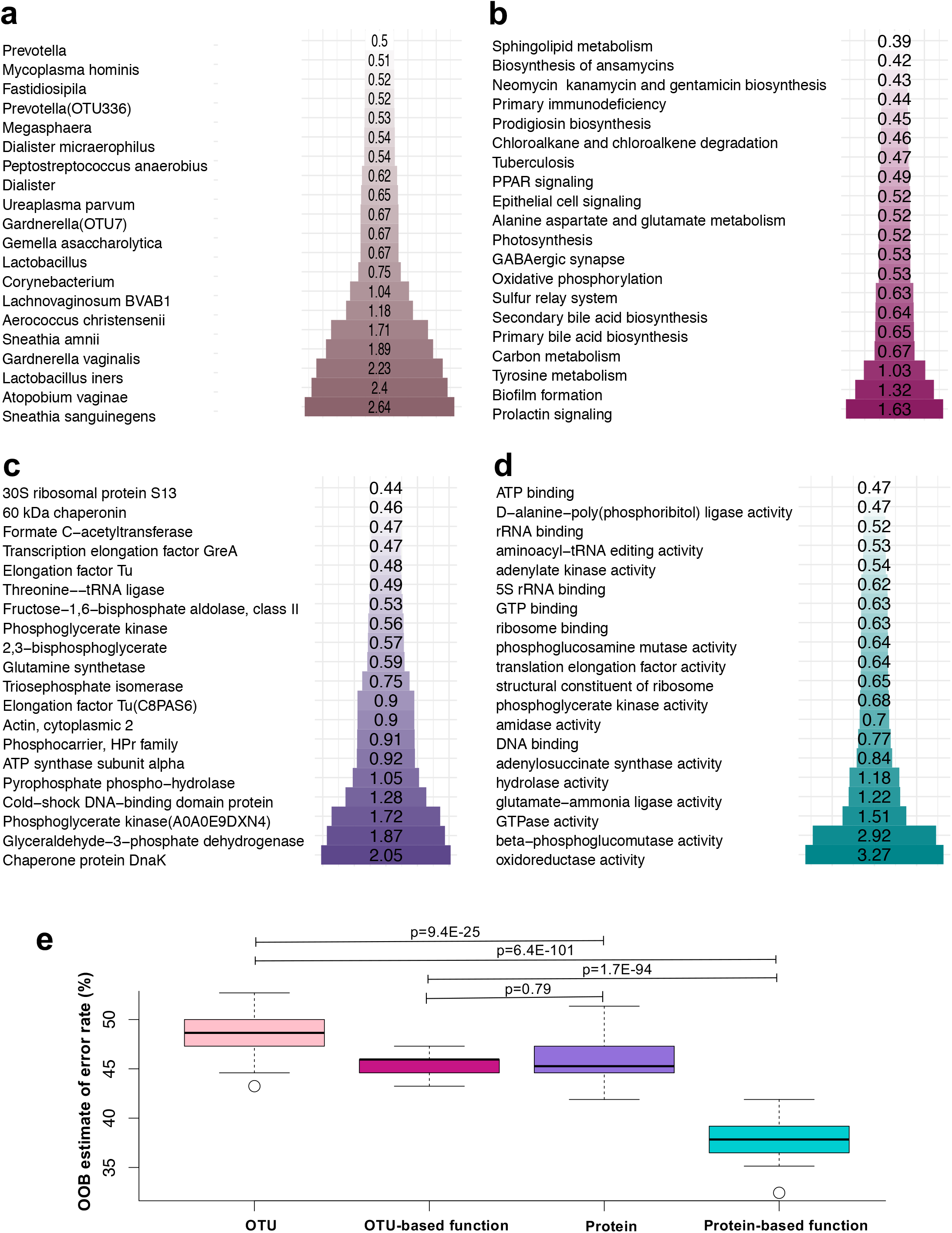
Comparison of bacterial, bacterial protein and bacterial function relative abundance for prediction of genital inflammation. Random forest analysis was used to evaluate the accuracy of (**a**) bacterial relative abundance (determined using 16S rRNA gene sequencing; n=74), (**b**) bacterial functional predictions based on 16 rRNA data (n=74), (**c**) bacterial protein relative abundance (determined using metaproteomics; n=74) and (**d**) bacterial protein molecular function relative abundance (determined by metaproteomics and aggregation of protein values assigned to the same gene ontology term; n=74) for determining the presence of genital inflammation (low, medium and high groups). Inflammation groups were defined based on hierarchical followed by K-means clustering of women according to the concentrations of nine pro-inflammatory cytokine concentrations [interleukin (IL)-1α, IL-1β, IL-6, IL-12p40, IL-12p70, tumor necrosis factor (TNF)-α, TNF-β, TNF-related apoptosis-inducing ligand (TRAIL), interferon (IFN)-γ]. The bars and numbers within the bars indicate the relative importance of each taxon, protein or function based on the Mean Decrease in Gini Value. The sizes of bars in each panel differ based on the length of the labels. (**e**) Each random forest model was iterated 100 times for each of the input datasets separately and the distribution of the out-of-bag error rates for the 100 models were then compared using t-tests. OTU: Operational taxonomic unit.

### Differences in microbial stress response, lactate dehydrogenase and metabolic pathways between women with high versus low FGT inflammation

Microbial functional profiles were further investigated by comparing molecular functions, biological processes and cellular component GO enrichment for detected proteins between women with low versus high inflammation using differential expression analysis, adjusting for potentially confounding variables including age, contraceptives, PSA and STIs (Fig. 4; Additional file 2: Table S5). The majority of microbial molecular functions (90/107), biological processes (57/83) and cellular components (14/23) were underrepresented in women with high compared to low genital inflammation, with most of the differentially abundant metabolic processes, including peptide, nucleoside, glutamine, and glycerophospholipid metabolic processes, decreased in women with inflammation. The phosphoenolpyruvate-dependent sugar phosphotransferase system (a major microbial carbohydrate uptake system including proteins assigned to *Lactobacillus* species, *S. amniii, A. vaginae*, and *G. vaginalis)* was also underrepresented in women with evidence of genital inflammation compared to women with low inflammation (FDR adj. p<0.0001; Fig. 4a). Additionally, L-lactate dehydrogenase activity was inversely associated with inflammation (Additional file 2: Table S5) and both large and small ribosomal subunit cellular components were underrepresented in those with high FGT inflammation (Fig. 4b). The only metabolic processes that were overabundant in women with high versus low inflammation were malate and ATP metabolic processes. Overrepresented microbial biological processes in women with genital inflammation also included response to oxidative stress and cellular components associated with cell division (e.g. mitotic spindle midzone) and ubiquitination (e.g. Doa10p ubiquitin ligase complex, Hrd1p ubiquitin ligase ERAD-L complex, RQC complex; Fig. 4).

**Figure 4.**
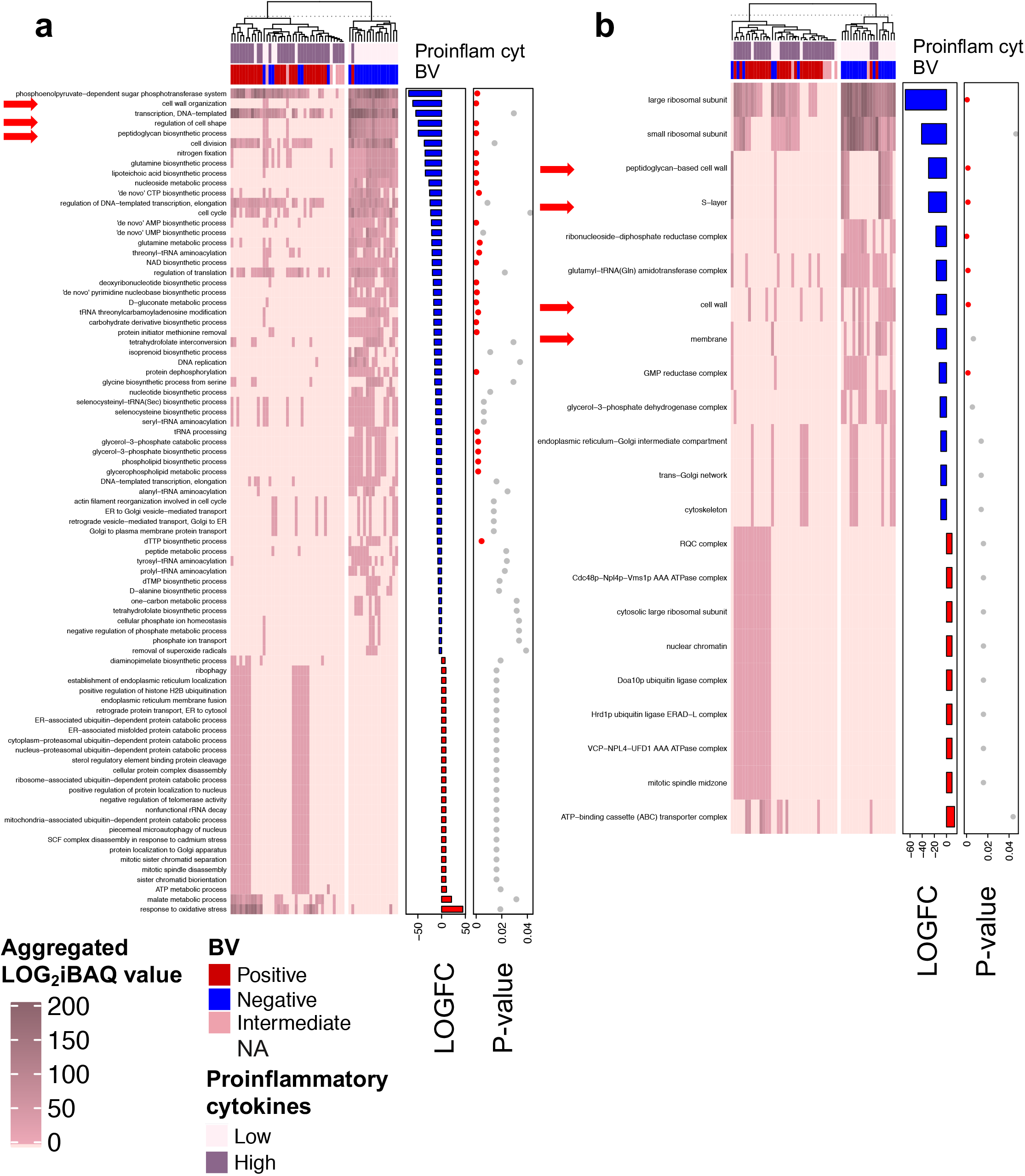
Microbial biological process and cellular component gene ontologies associated with genital inflammatory cytokine profiles. Unsupervised hierarchical clustering of aggregated intensity-based absolute quantification (iBAQ) values for microbial protein (**a**) biological process (BP) or (**b**) cellular component (CC) gene ontology (GO) relative abundance (n=113). GO relative abundance was determined by metaproteomics and aggregation of microbial protein iBAQ values assigned to the same GO term. The heatmaps show aggregated microbial GO relative abundance and the bar graphs show fold changes in aggregated log_2_-transformed iBAQ values (LOGFC) for each microbial protein with the same (**a**) BP or (**b**) CC GO in women with high versus low inflammation. Inflammation groups were defined based on hierarchical followed by K-means clustering of women according to the concentrations of nine pro-inflammatory cytokine concentrations [interleukin (IL)-1α, IL-1β, IL-6, IL-12p40, IL-12p70, tumor necrosis factor (TNF)-α, TNF-β, TNF-related apoptosis-inducing ligand (TRAIL), interferon (IFN)-γ]. Red bars indicate positive and blue bars indicate negative fold changes in women with high versus low inflammation. False discovery rate adjusted p-values are shown by dots, with red dots indicating low p-values. Red arrows indicate cell wall and membrane processes and components. BV: Bacterial vaginosis. Proinflam cyt: Pro-inflammatory cytokine

### Underrepresented *Lactobacillus* peptidoglycan, cell wall and membrane pathways in women with high versus low FGT inflammation

Among the top biological processes inversely associated with inflammation were cell wall organization, regulation of cell shape and peptidoglycan biosynthetic process (all adj. p≤0.0001; Fig. 4a). The GO terms, regulation of cell shape, and peptidoglycan biosynthetic process included identical proteins (e.g. D-alanylalanine synthetase, D-alanyl-D-alanine-adding enzyme and Bifunctional protein GlmU), while cell wall organization included the same proteins plus D-alanyl carrier protein. Although proteins involved in these processes were only detected for *Lactobacillus* species and may thus be biomarkers of the increased relative abundance of lactobacilli, each remained significantly associated with inflammation after adjusting for the relative abundance of *L. iners* and *non-iners Lactobacillus* species (determined using 16S rRNA gene sequencing), as well as STI status, PSA detection and hormonal contraceptive use [beta-coefficient: −1.68; 95% confidence interval (CI): −2.84 to −0.52; p=0.004 for cell wall organization; beta-coefficient: −1.71; 95% CI: −2.91 to −0.53; p=0.005 for both regulation of cell shape and peptidoglycan biosynthetic process]. This suggests that the relationship between these processes and inflammation is independent of the relative abundance of *Lactobacillus* species and that the cell wall and membrane properties of lactobacilli may play a role in modulating FGT inflammation. Furthermore, we found that multiple cell wall and membrane cellular components, including the S-layer, peptidoglycan-based cell wall, cell wall and membrane, were similarly underrepresented in women with high versus low FGT inflammation (Fig. 4b). Using metaproteomic data from an independent group of 10 women, we similarly found that peptidoglycan biosynthetic process, cell wall organization, and regulation of cell shape GOs were underrepresented in women with high inflammation (Fig. 5a).

**Figure 5.**
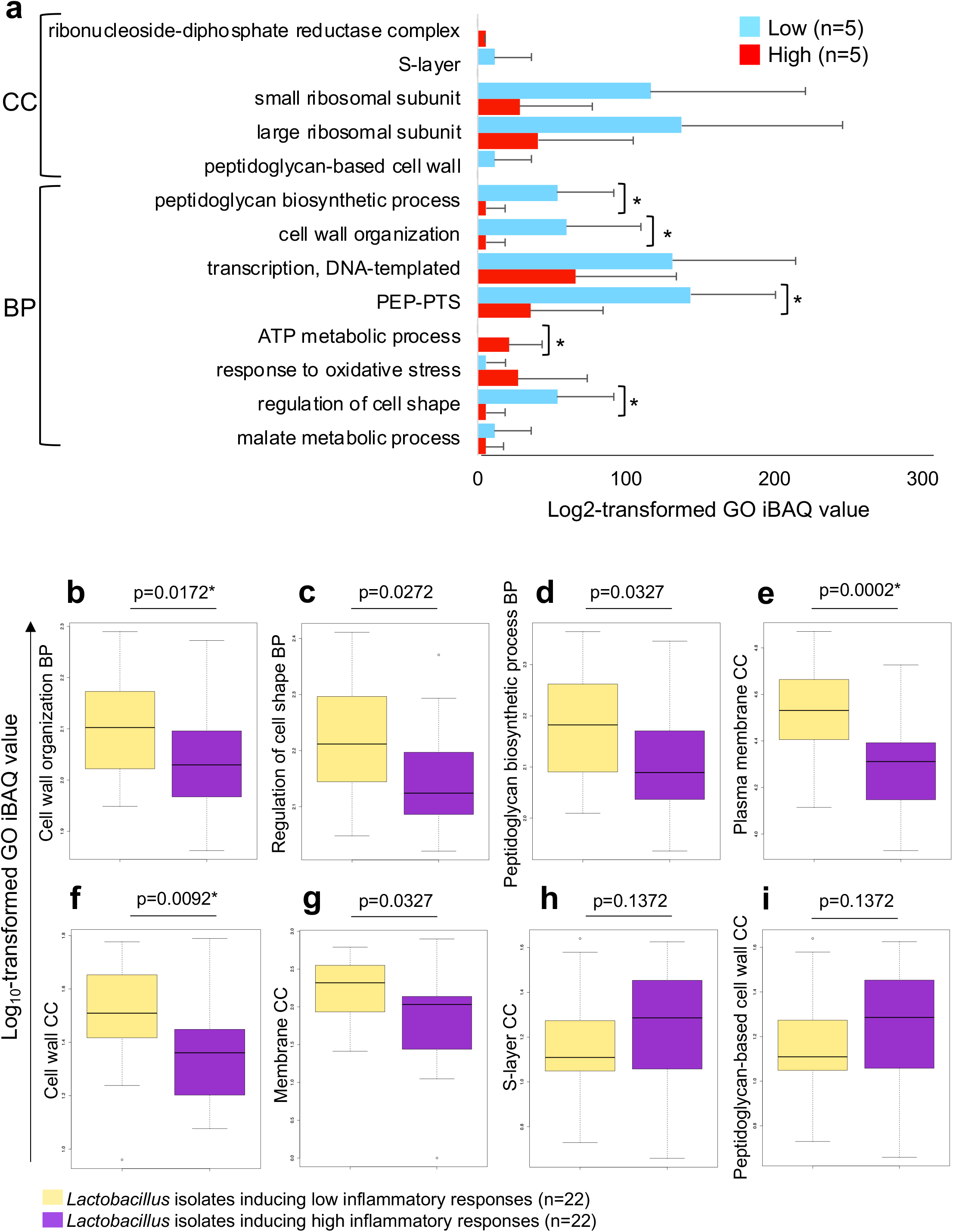
Relative abundance of gene ontologies in independent samples and in inflammatory versus non-inflammatory *Lactobacillus* isolates. **(a)** The top 14 microbial biological process (BP) and cellular component (CC) gene ontology (GO) terms that distinguished women with low versus high inflammation in the full cohort (n=113) were validated in an independent sample of ten women from Cape Town, South Africa. Liquid chromatography-tandem mass spectrometry was used to evaluate the metaproteome in lateral vaginal wall swabs from these women. Inflammation groups were defined based on hierarchical followed by K-means clustering of these ten women according to the concentrations of nine pro-inflammatory cytokine concentrations [interleukin (IL)-1α, IL-1β, IL-6, IL-12p40, IL-12p70, tumor necrosis factor (TNF)-α, TNF-β, TNF-related apoptosis-inducing ligand (TRAIL), interferon (IFN)-γ]. GO relative abundance was determined by aggregation of microbial protein intensity-based absolute quantification (iBAQ) values assigned to a same GO term. The relative abundance of the top BPs and CCs (except ATP binding cassette transporter complex which was not detected in these samples) are shown as bar graphs, with blue indicating women with low inflammation (n=5) and red indicating women with high inflammation (n=5). Welch’s t-test was used for comparisons. *P<0.05. **(b-i)** Twenty-two *Lactobacillus* isolates that induced relatively high inflammatory responses and 22 isolates that induced lower inflammatory responses were adjusted to 4.18 x 10^6^ CFU/ml in bacterial culture medium and incubated for 24 hours under anaerobic conditions. Following incubation, protein was extracted, digested and liquid chromatography-tandem mass spectrometry was conducted. Raw files were processed with MaxQuant against a database including the *Lactobacillus* genus and common contaminants. The iBAQ values for proteins with the same gene ontologies were aggregated, log_10_-transformed and compared using Mann-Whitney U test. Box-and-whisker plots show log_10_-transformed iBAQ values, with lines indicating medians and whiskers extending to 1.5 times the interquartile range from the box. A false discovery rate step-down procedure was used to adjust for multiple comparisons and adjusted p-values <0.05 were considered statistically significant.

To further investigate whether the associations between FGT inflammation and *Lactobacillus* biological processes and cellular components were independent of *Lactobacillus* relative abundance, we compared these GOs in BV negative women with *Lactobacillus* dominant communities who were grouped (median splitting of samples based on the first principal component of nine pro-inflammatory cytokine concentrations) as having low (n=20) versus high (n=20) inflammation (Fig. S6). Interestingly, it was found that, even though only non-significant trends towards decreased relative abundance of lactobacilli determined by 16S rRNA gene sequencing (p=0.73) and metaproteomics (p=0.14) were observed, several previously identified GOs, including peptidoglycan-based cell wall (p=0.01), cell wall organization (p=0.03), and S-layer (p=0.01), remained associated with the level of inflammation (Fig. S6). This suggests an association between these biological processes and cellular components that is not fully explained by the relative abundance of the lactobacilli themselves.

### *In vitro* confirmation of the role of *Lactobacillus* cell wall and membrane pathways in regulating genital inflammation

As previous studies have suggested that the cell wall, as well as the cell membrane, may influence the immunomodulatory properties of *Lactobacillus* species and that peptidoglycan may be directly immunosuppressive [14,24], we further investigated the relationships between *Lactobacillus* cell wall and membrane properties and inflammation using 64 *Lactobacillus* species that were isolated from the same women [25]. From these lactobacilli, the 22 isolates that induced the lowest levels of pro-inflammatory cytokine production by vaginal epithelial cells and the 22 isolates that induced the greatest cytokine responses were selected and their protein profiles evaluated using LC-MS/MS. Cell wall organization, regulation of cell shape, and peptidoglycan biosynthetic process that were associated with genital inflammation in the FGT metaproteomic analysis of vaginal swab samples were also significantly overrepresented in isolates that induced low compared to high levels of inflammatory cytokines *in vitro* (Fig. 5b-d). Furthermore, plasma membrane, cell wall and membrane cellular components were similarly overrepresented in isolates that induced low versus high levels of inflammation (Fig. 5e-i). The cell wall cellular component included multiple proteins involved in adhesion (adhesion exoprotein, LPXTG-motif cell wall anchor domain protein, collagen-binding protein, mannose-specific adhesin, putative mucin binding protein, sortase-anchored surface protein and surface anchor protein). The relative abundance of this cell wall cellular component correlated positively with the level of adhesion of the isolates to vaginal epithelial cells *in vitro* (rho=0.30, p=0.0476), which in turn correlated inversely with *in vitro* inflammatory cytokine production by these cells in response to the isolates [25]. Importantly, the cell wall organization biological process included multiple proteins involved in peptidoglycan biosynthesis, which has the potential to dampen immune responses in a strain-dependent manner [14,26].

### Changes in FGT metaproteomic profiles over time

To evaluate variations in FGT metaproteomic profiles associated with inflammatory profiles over time, we compared proteins, taxa and biological process and cellular component GOs between two time-points nine weeks apart (n=74). The grouping according to inflammatory cytokine profile was consistent at both visits for 41/74 (55%) participants, including 7/12 (58%) women who received antibiotic treatment between visits (Fig. 6a). The findings based on the second visit also confirmed the role of inflammation as one of the drivers of variation in complex metaproteome data, and similar patterns of data distribution according to inflammation status were observed at both visits (Fig. 6b). When the proteins, taxa and functions that were most closely associated with inflammation grouping (high vs. low) at the first visit, were evaluated at the second visit, overall profiles remained similar between visits (Fig. 6c-e). Significantly lower relative abundance of *Lactobacillus* species and increased non-optimal bacteria were observed in women with inflammation at the second visit (Fig. 6d). Similarly, cell wall organization, regulation of cell shape, and peptidoglycan biosynthetic process were among the top biological processes associated with low versus high inflammation at both visits (for other top GO terms see Fig. 6e). Among women assigned to the same inflammatory profile group at both visits, the relative abundance of cell wall-related GO terms, malate metabolic process and response to oxidative stress were highly consistent (Additional file 1: Fig. S7a-c). However, decreased levels of inflammation between visits were associated with an increase in the relative abundance of cell wall GOs and decreased response to oxidative stress and malate metabolic process GOs (Additional file 1: Fig. S7d-f). On the other hand, increased levels of inflammation were associated with decreased relative abundance of cell wall GOs and increased response to oxidative stress and malate metabolic process GOs (Additional file’: Fig. S7g-i).

**Figure 6.**
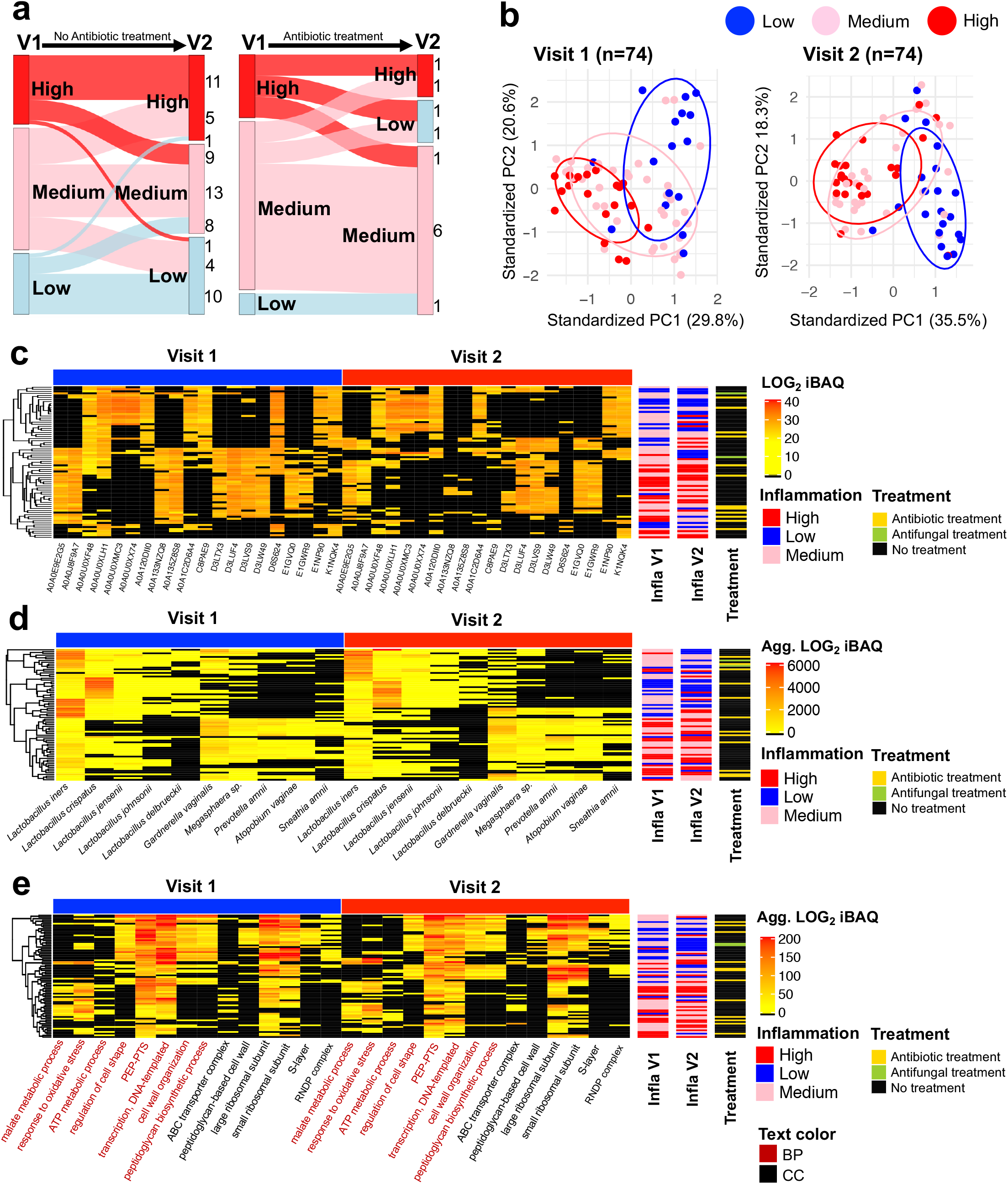
Longitudinal changes in FGT metaproteomic profiles. Liquid chromatography-tandem mass spectrometry was used to evaluate the metaproteome in lateral vaginal wall swabs from 74 women from Cape Town, South Africa at two visits nine weeks (interquartile range: 9-11 weeks) apart. (**a**) Sankey diagram was used to visualize changes in inflammatory cytokine profiles between visits. (**b**) Principal component analysis (mixOmics R package) was used to group women based on the log_2_-transformed intensity-based absolute quantification (iBAQ) values of all proteins identified at both visits. Grouping is based on vaginal pro-inflammatory cytokine concentrations and each point represents an individual woman. (**c**) Top 20 proteins (UniProt IDs) distinguishing women with and without inflammation at both visits determined by moderated t-test (limma R package) and random forest analysis (randomForest R package). (**d**) Top 10 taxa distinguishing women with and without inflammation at both visits determined by moderated t-test (limma R package) and random forest algorithm (randomForest R package). (**e**) Top 14 biological process and cellular component gene ontology terms distinguishing women with and without inflammation at both visits determined by moderated t-test (limma R package) and random forest algorithm (randomForest R package). Positions of participants in each heatmap are fixed and the inflammation status of each participant across the visits can be tracked using the row sidebars. Inflammation groups were defined based on hierarchical followed by K-means clustering of nine pro-inflammatory cytokine concentrations [interleukin (IL)-1α, IL-1β, IL-6, IL-12p40, IL-12p70, tumor necrosis factor (TNF)-α, TNF-β, TNF-related apoptosis-inducing ligand (TRAIL), interferon (IFN)-γ]. ABC: ATP-binding cassette. Agg: aggregated. BP: biological process. CC: cellular component. Infla: inflammation. PC: principal component. PEP-PTS: phosphoenolpyruvate-dependent phosphotransferase system. RNDP complex: ribonucleoside-diphosphate reductase complex.

## Discussion

Understanding biological risk factors for HIV acquisition is critical for the development of effective HIV prevention strategies. FGT inflammation, defined by elevated inflammatory cytokine levels, has been identified as a key factor associated with HIV infection risk. However, the causes and underlying mechanisms are not fully understood. In this study, we demonstrate that, while the presence of particular microorganisms, including non-optimal bacteria and *Lactobacillus* species, is important for modulating FGT inflammation, their activities and functions are likely of greater importance. We showed that the molecular functions of bacterial proteins predicted FGT inflammation status more accurately than taxa relative abundance determined using 16S rRNA gene sequencing, as well as functional predictions based on 16S rRNA gene sequencing analysis. The majority of microbial biological processes were underrepresented in women with high compared to low inflammation (57/83), including metabolic pathways and a strong signature of reduced *Lactobacillus-associated* cell wall organization and peptidoglycan biosynthesis. This signature remained associated with FGT inflammation after adjusting for the relative abundance of lactobacilli, as well as other potential confounders (such as hormone contraceptive use, semen exposure, STIs), and was also associated with inflammatory cytokine induction in vaginal epithelial cells in response to *Lactobacillus* isolates *in vitro*. We observed a high level of consistency between FGT metaproteomics profiles of the same women at two time points nine weeks apart, particularly among those with the same level of inflammation at both visits. However, cell wall organization and peptidoglycan biosynthesis increased between visits in women who moved from a higher to a lower inflammation grouping and decreased in women who moved from a lower to a higher inflammation grouping.

While we found a large degree of similarity between metaproteomics and 16S rRNA gene sequence taxonomic assignment determined for the same women [27], important differences in the predicted relative abundance of most bacteria were observed. Differences in the relative abundance of certain taxa observed using metaproteomics and 16S rRNA gene sequence data is not surprising as taxonomic assignment from metaproteomic approaches also indicates functional activity (measured by the amount of expressed protein), rather than simply the abundance of a particular bacterial taxon. Additionally, others have noted that 16S rRNA gene sequencing does not have the ability to distinguish between live bacteria and transient DNA [28]. We also observed differences that are likely due to database limitations. For example, *Lachnovaginosum* genomospecies (previously known as BVAB1) was not identified in the metaproteomics analysis. This is due to the fact that this species was not present in the UniProt or NCBI databases [29]. However, the *Clostridium* and *Ruminococcus* species which were identified are likely to be *Lachnovaginosum* genomospecies since these taxa fall into the same Clostridiales order. Additionally, *Clostridium* and *Ruminococcus* species were more abundant in women with high compared to low inflammation, as expected for *Lachnovaginosum* genomospecies [10]. Metaproteomic analysis was able to identify lactobacilli to species level, while sequencing of the V4 region of the 16S rRNA gene had more limited resolution for this genus, as expected.

The finding that fungal protein relative abundance was substantially lower than bacterial protein relative abundance is not surprising as it has been estimated that the total number of fungal cells is orders of magnitude lower than the number of bacterial cells in the human body [30]. Although *Candida* proteins were detected, the curated dataset (following removal of taxa that had only 1 protein detected or 2 proteins detected in only a single sample) did not include any *Candida* species. This may be due to the relative rarity of fungal cells in these samples and it would be interesting to evaluate alternative methodology for sample processing to better resolve this population in the future [31]. A limitation of microbial functional analysis using metaproteomics is that, at present, there is generally sparse population of functional information in the databases for the majority of the microbiome. Thus, the findings of this study will be biased toward the microbes that are better annotated. Nonetheless, microbial function was closely associated with genital inflammatory profiles and it is possible that this relationship may be even stronger should better annotation exist. Another limitation of this study is that the *in vitro Lactobacillus* analysis was conducted using a transformed cell line system that is a greatly simplified model and cannot recapitulate the complexity of the FGT and the microbiome. It is however significant to note that similar proteomic signatures were noted both *in vitro* and *in vivo*, regardless of this limitation. To enable researchers to further study associations between microbial composition and function with BV, STIs, and FGT inflammation by conducting detailed analysis of specific proteins, taxa and GO terms of interest, we have developed the “FGT-METAP” online application. The application is available at (http://immunodb.org/FGTMetap/) and allows users to repeat the analyses presented here, as well as obtain additional information by analyzing individual proteins, species, and pathways.

A significant functional signature associated with FGT inflammation grouping included *Lactobacillus-associated* cell wall, peptidoglycan and cell membrane biological processes and cellular components. Although these functions were linked exclusively to *Lactobacillus* species, the associations with FGT inflammation were upheld after adjusting for *Lactobacillus* relative abundance determined by 16S rRNA gene sequencing and were also evident in an analysis including only women with *Lactobacillus*-dominant microbiota. This profile was confirmed when these GOs were compared between *Lactobacillus* isolates that induced high versus low levels of cytokine production *in vitro*. Previous studies have suggested that the peptidoglycan structure, the proteins present in the cell wall, as well as the cell membrane, may influence the immunomodulatory properties of *Lactobacillus* species [14,24]. In a murine model, administration of peptidoglycan extracted from gut lactobacilli was able to rescue the mice from colitis [26]. This activity was dependent on the NOD2 pathway and correlated with an upregulation of the indoleamine 2,3-dioxygenase immunosuppressive pathway. The immunomodulatory properties of peptidoglycan were furthermore dependent on the strain of lactobacilli from which the peptidoglycan was isolated [26]. The increase in cell wall organization and peptidoglycan biosynthesis that accompanied a reduction in local inflammation suggests an increase in *Lactobacillus* strains producing relatively large amounts of proteins involved in these processes. In a previous study, we showed that adhesion of lactobacilli to vaginal epithelial cells *in vitro* was inversely associated with cytokine responses, suggesting that direct interaction between the isolates and vaginal epithelial cells is critical for immunoregulation [25]. In addition to cell wall and membrane properties, L-lactate dehydrogenase activity was identified as a molecular function inversely associated with inflammation. Similarly, D-lactate dehydrogenase relative abundance and D-lactate production by the *Lactobacillus* isolates included in this study were also associated with inflammatory profiles in an *in vitro* analysis [25]. These findings support the results of previous studies showing that lactic acid has immunoregulatory properties [16]. Taken together, these findings suggest that lactobacilli may modulate the inflammatory environment in the FGT through multiple mechanisms. It was further found that *L. crispatus* was more strongly associated with low inflammation in the differential abundance analysis and that *L. iners* was the *Lactobacillus* sp. most frequently detected in women with high inflammation. The co-expression analysis revealed that the majority of *L. crispatus* proteins grouped separately to *L. iners* proteins. However, both the *L. crispatus* and the *L. iners* modules were strongly inversely associated with inflammatory cytokine and chemokine concentrations. This is interesting since *L. crispatus* dominance is considered optimal, while the role of *L. iners*, the most prevalent *Lactobacillus* species in African women [10,11], is poorly understood and it has been associated with compositional instability and transition to a non-optimal microbiota, as well as increased risk of STI acquisition [32,33].

In this study, we observed a similar host proteome profile associated with high FGT inflammation compared to previous studies [7]. Multiple inflammatory pathways were overrepresented in women with high versus low FGT inflammation, while signatures of reduced barrier function were observed in women with high inflammation, with underabundant endothelial, ectoderm and tight junction biological processes. These findings similarly suggest that genital inflammation may be associated with epithelial barrier function.

## Conclusions

The link between FGT microbial function and local inflammatory responses described in this study suggests that both the presence of specific microbial taxa in the FGT and their properties and activities likely play a critical role in modulating inflammation. Currently, the annotation of microbial functions is sparse, but with ever-increasing amounts of high-quality, high-throughput data, the available information will improve steadily. The findings of the present study contribute to our understanding of the mechanisms by which the microbiota may influence local immunity, and in turn alter the risk of HIV infection. Additionally, the analyses described herein identify specific microbial properties that may be harnessed for biotherapeutic development.

## Methods

### Participants

This study included sexually experienced HIV-negative adolescent girls and young women (16-22 years) from the Women’s Initiative in Sexual Health (WISH) study in Cape Town, South Africa [22]. While the parent study cohort comprised 149 women, the present sub-study included 113 women who had both vaginal swabs available for metaproteomics analysis and menstrual cup (MC) cervicovaginal secretions available for cytokine profiling. Seventy-four of these women had specimens available at a second time-point 9 weeks later (interquartile range: 9-11 weeks) for metaproteomics analysis. Of these women, 12 received antibiotic treatment for laboratory-diagnosed or symptomatic STIs and/or BV (doxycycline, metronidazole, azithromycin, cefixime, ceftriaxone and/or amoxicillin) between study visits, while 2 received topical antifungal treatment (clotrimazole vaginal cream). Ten additional women who did not have visit 1 samples available for analysis were included for validation purposes. The University of Cape Town Faculty of Health Sciences Human Research Ethics Committee (HREC REF: 267/2013) approved this study. Women ≥18 years provided written informed consent, those <18 years provided informed assent and informed consent was obtained from parents/guardians.

### Definition of genital inflammation based on cytokine concentrations

For cytokine measurement, MCs (Softcup^®^, Evofem Inc, USA) were inserted by the clinician and kept in place for an hour, after which they were transported to the lab at 4°C and processed within 4 hours of removal from the participants. MCs were centrifuged and the cervicovaginal secretions were re-suspended in phosphate buffered saline (PBS) at a ratio of 1ml of mucus: 4 ml of PBS and stored at −80°C until cytokine measurement. Prior to cytokine measurement, MC secretions were pre-filtered using 0.2 μm cellulose acetate filters (Sigma-Aldrich, MO, USA). The concentrations of 48 cytokines were measured in MC samples using Luminex (Bio-Rad Laboratories Inc^®^, CA, USA) [23]. K-means clustering was used to identify women with low, medium and high pro-inflammatory cytokine [interleukin (IL)-1α, IL-1β, IL-6, IL-12p40, IL-12p70, tumor necrosis factor (TNF)-α, TNF-β, TNF-related apoptosis-inducing ligand (TRAIL), interferon (IFN)-γ] and chemokine profiles [cutaneous T-cell-attracting chemokine (CTACK), eotaxin, growth regulated oncogene (GRO)-α, IL-8, IL-16, IFN-γ-induced protein (IP)-10, monocyte chemoattractant protein (MCP)-1, MCP-3, monokine induced by IFN-γ (MIG), macrophage inflammatory protein (MIP)-1α, MIP-1β, regulated on activation, normal T cell expressed and secreted (RANTES)] (Additional file 1: Fig. S2).

### STI and BV diagnosis and evaluation of semen contamination

Vulvovaginal swab samples were screened for common STIs, including *Chlamydia trachomatis, Neisseria gonorrhoeae, Trichomonas vaginalis, Mycoplasma genitalium*, herpes simplex virus (HSV) types 1 and 2, *Treponema pallidum* and *Haemophilus ducreyi*, using real-time multiplex PCR assays [22]. Lateral vaginal wall swabs were collected for BV assessment by Nugent scoring. Blood was collected for HIV rapid testing (Alere Determine™ HIV-1/2 Ag/Ab Combo, Alere, USA). PSA was measured in lateral vaginal wall swabs using Human Kallikrein 3/PSA Quantikine ELISA kits (R&D Systems, MN, USA).

### Discovery metaproteomics and analysis

For shotgun LC-MS/MS, lateral vaginal wall swabs were collected, placed in 1 ml PBS, transported to the laboratory at 4°C and stored at −80°C. Samples were computationally randomized for processing and analysis. The stored swabs were thawed overnight at 4°C before each swab sample was vortexed for 30 seconds, the mucus scraped off the swab on the side of the tube and vortexed for an additional 10 seconds. The samples were then clarified by centrifugation and the protein content of each supernatant was determined using the Quanti-Pro bicinchoninic acid (BCA) assay kit (Sigma-Aldrich, MO, USA). Equal protein amounts (100 μg) were denatured with urea exchange buffer, filtered, reduced with dithiothreitol, alkylated with iodoacetamide and washed with hydroxyethyl piperazineethanesulfonic acid (HEPES). Acetone precipitation/formic acid (FA; Sigma-Aldrich, MO, USA) solubilisation was added as a further sample clean-up procedure. The samples were then incubated overnight with trypsin (Promega, WI, USA) and the peptides were eluted with HEPES and dried via vacuum centrifugation. Reversed-phase liquid chromatography was used for desalting using a step-function gradient. The eluted fractions were dried via vacuum centrifugation and kept at −80°C until analysis. LC-MS/MS analysis was conducted on a Q-Exactive quadrupole-Orbitrap MS (Thermo Fisher Scientific, MA, USA) coupled with a Dionex UltiMate 3000 nano-UPLC system (120min per sample). The solvent system included Solvent A: 0.1% Formic Acid (FA) in LC grade water (Burdick and Jackson, NJ, USA); and Solvent B: 0.1% FA in Acetonitrile (ACN; Burdick & Jackson, NJ, USA).

The MS was operated in positive ion mode with a capillary temperature of 320°C, and 1.95 kV electrospray voltage was applied. All 187 samples were evaluated in 11 batches over a period of 3 months. A quality control consisting of pooled lateral vaginal swab eluants from the study participants was run at least twice with every batch. Preliminary quality control analysis was conducted, and flagged samples/batches were rerun.

Due to the important role of database in metaproteomic analysis, we compared two different databases to identify proteins in our dataset: (i) UniProt database restricted to human and microbial entries (73,910,451, release August-2017) and filtered using the MetaNovo pipeline [34]; (ii) vaginal metagenome-based database with human proteins obtained from the study of Afiuni-Zadeh et al. [35]. The raw data were processed using MaxQuant version 1.5.7.4 against each database, separately. The detailed parameters are provided in Additional file 1: Table S6. In brief, methionine oxidation and acetylation of the protein N-terminal amino acid were considered as variable modifications and carbamidomethyl (C) as a fixed modification. The digestion enzyme was trypsin with a maximum of two missed cleavages. The taxonomic analysis of the proteins identified using both of the databases was comparable, however, the first database (filtered UniProt human and microbial entries) resulted in the identification of a greater number of proteins compared to the vaginal metagenome database, whilst exhibiting a high degree of similarity compared to the 16S rRNA gene sequence data. Therefore, the data generated using the first database was used for downstream analysis.

For quality control analysis, raw protein intensities and log_10_-transformed iBAQ intensities of the quality control pool, as well as the raw protein intensities of the clinical samples, were compared between batches (not shown). As variation in raw intensity was observed, all data were adjusted for batch number prior to downstream analysis. The cleaned and transformed iBAQ values were used to identify significantly differentially abundant proteins according to inflammatory cytokine profiles, BV status or STIs using the limma R package [36]. The p-values were obtained from the moderated t-test after adjusting for confounding variables including age, contraceptives, PSA, and STIs. Proteins with FDR adjusted p-values ≤0.05 and log_2_-transformed fold change ≥ 1.2 or ≤ −1.2 were considered as significantly differentially abundant. To define significantly differentially abundant taxa and GOs, aggregated iBAQ values of proteins of the same species or GO IDs were compared between women with low, medium and high inflammation using the limma R package. The aggregation was performed separately for host and microbial proteins. The p-values along with GO IDs were uploaded to REVIGO (http://revigo.irb.hr/) to visualize the top 50 biological processes with the lowest p-values.

The effects of BV, STIs and chemokine and pro-inflammatory cytokine profiles on the metaproteome were investigated by PCA using the mixOmics R package [37]. The ComplexHeatmap package [38] was used to generate heatmaps and cluster the samples and metaproteomes based on the hierarchical and k-means clustering methods. Additionally, R packages EnhancedVolcano, weighted correlation network analysis (WGCNA) [39], and FlipPlots were applied for plotting volcano plots, co-correlation analysis, and Sankey diagrams, respectively. Basic R functions and the ggplot2 R package were used for data manipulation, transformation, normalization and generation of graphics. To identify the key factors distinguishing women defined as having low, medium and high inflammation in their FGTs, we used random forest analysis using the R package randomForest [40] using the following settings: (i) the type of random forest was classification, (ii) the number of trees was 2000, and (iii) the number of variables evaluated at each split was one-third of the total number of variables [proteins/molecular function GO IDs/16S rRNA gene-based operational taxonomic units (OTUs)]. To compare the accuracy of the protein-, function- and species-based models, we iterated each random forest model 100 times for each of the input datasets separately. The distributions of the OOBs for 100 models for each data type were then compared using t-tests. For the WGCNA analysis, a microbial-host weighted co-correlation network was built using iBAQ values for all proteins and modules were identified using dynamic tree cut. Correlations between proteins and modules were then determined and the top taxa and biological processes assigned to each of the proteins identified. Spearman’s rank correlation was used to determine correlation coefficients between individual cytokines/chemokines and module eigengenes (the first principal component of the expression matrix of the corresponding module) for all samples. Average Spearman’s correlation coefficients for all pro-inflammatory cytokines and chemokines were then determined. Similarly, Spearman’s correlation coefficients were calculated between module eigengenes and Nugent scores and STIs (including *C. trachomatis, N. gonorrhoeae, T. vaginalis, M. genitalium* and active HSV-2).

### 16S rRNA gene sequencing and analysis

Bacterial 16S rRNA gene sequencing and analysis was conducted as previously described [27]. Briefly, the V4 hypervariable region of the 16S rRNA gene was amplified using modified universal primers [41]. Duplicate samples were pooled, purified using Agencourt AMPure XP beads (Beckman Coulter, CA, USA) and quantified using the Qubit dsDNA HS Assay (Life Technologies, CA, USA). Illumina sequencing adapters and dual-index barcodes were added to the purified amplicon products using limited cycle PCR and the Nextera XT Index Kit (Illumina, CA, USA). Amplicons from 96 samples and controls were pooled in equimolar amounts and the resultant libraries were purified by gel extraction and quantified (Qiagen, Germany). The libraries were sequenced on an Illumina MiSeq platform (300 bp paired-end with v3 chemistry). Following de-multiplexing, raw reads were preprocessed, merged and filtered using DADA2 [42]. Primer sequences were removed using a custom python script and the reads truncated at 250 bp. Taxonomic annotation was based on a customized version of SILVA reference database. Only samples with ≥ 2000 reads were included in further analyses. The representative sequences for each amplicon sequence variant were BLAST searched against NCBI non-redundant protein and 16S rRNA gene sequence databases for further validation, as well as enrichment of the taxonomic classifications. Microbial functional predictions were performed using Piphillin server [43] against the Kyoto Encyclopedia of Genes and Genomes (KEGG) database (99% identity cutoff). The relative abundance of functional pathways (KEGG level 3) was used for RF analysis.

### *Lactobacillus* isolation

Lactobacilli were isolated from cervicovaginal secretions collected using MCs by culturing in de Man Rogosa and Sharpe (MRS) broth for 48 hours at 37°C under anaerobic conditions. The cultures were streaked onto MRS agar plates under the same culture conditions, single colonies were picked and then pre-screened microscopically by Gram staining. Matrix Assisted Laser Desorption Ionization Time of Flight (MALDI-TOF) was conducted to identify the bacteria to species level. A total of 115 lactobacilli were isolated and stored at −80°C in a final concentration of 60% glycerol. From these, 64 isolates from 25 of the study participants were selected for evaluation of inflammatory profiles.

### Vaginal epithelial cell stimulation and measurement of cytokine concentrations

As described previously [25], vaginal epithelial cells (VK2/E6E7 ATCC^®^ CRL-2616™; RRID:CVCL_6471) were maintained in complete keratinocyte serum free media (KSFM) supplemented with 0.4 mM calcium chloride, 0.05 mg/ml of bovine pituitary extract, 0.1 ng/ml human recombinant epithelial growth factor and 50 U/ml penicillin and 50 U/ml streptomycin (Sigma-Aldrich, MO, USA). The VK2 cells were seeded into 24-well tissue culture plates, incubated at 37°C in the presence of 5% carbon dioxide and grown to confluency. Lactobacilli, adjusted to 4.18 x 10^6^ colony forming units (CFU)/ml in antibiotic-free KSFM, were used to stimulate VK2 cells for 24 hours at 37°C in the presence of 5% carbon dioxide. As previously described, *G. vaginalis* ATCC 14018 cultures standardized to 1 x 10^7^ CFU/ml in antibiotic-free KSFM were used as a positive control [25]. Production of IL-6, IL-8, IL-1α, IL-1β, IP-10, MIP-3α, MIP-1α, MIP-1β and IL-1RA were measured using a Magnetic Luminex Screening Assay kit (R&D Systems, MN, USA) and a Bio-Plex™ Suspension Array Reader (Bio-Rad Laboratories Inc^®^, CA, USA). VK2 cell viability following bacterial stimulation was confirmed using the Trypan blue exclusion assay. PCA was used to compare the overall inflammatory profiles of the isolates and to select the 22 most inflammatory and 22 least inflammatory isolates for characterization using proteomics.

### Characterization of *Lactobacillus* protein expression *in vitro* using mass spectrometry

Forty-four *Lactobacillus* isolates were adjusted to 4.18 x 10^6^ CFU/ml in MRS and incubated for 24 hours under anaerobic conditions. Following incubation, the cultures were centrifuged and the pellets washed 3x with PBS. Protein was extracted by resuspending the pellets in 100 mM triethylammonium bicarbonate (TEAB; Sigma-Aldrich, MO, USA) 4% sodium dodecyl sulfate (SDS; Sigma-Aldrich, MO, USA). Samples were sonicated in a sonicating water bath and subsequently incubated at 95°C for 10min. Nucleic acids were degraded using benzonase nuclease (Sigma-Aldrich, MO, USA) and samples were clarified by centrifugation at 10 000xg for 10 min. Quantification was performed using the Quanti-Pro BCA assay kit (Sigma-Aldrich, MO, USA). HILIC beads (ReSyn Biosciences, South Africa) were washed with 250 μl wash buffer [15% ACN, 100 mM Ammonium acetate (Sigma-Aldrich, MO, USA) pH 4.5]. The beads were then resuspended in loading buffer (30% ACN, 200mM Ammonium acetate pH 4.5). A total of 50 μg of protein from each sample was transferred to a protein LoBind plate (Merck, NJ, USA). Protein was reduced with tris (2-carboxyethyl) phosphine (Sigma-Aldrich, MO, USA) and alkylated with methylmethanethiosulphonate (MMTS; Sigma-Aldrich, MO, USA). HILIC magnetic beads were added at an equal volume to that of the sample and a ratio of 5:1 total protein and incubated on the shaker at 900 rpm for 30 min. After binding, the beads were washed four times with 95% ACN. Protein was digested by incubation with trypsin for four hours and the supernatant containing peptides was removed and dried down in a vacuum centrifuge. LC-MS/MS analysis was conducted with a Q-Exactive quadrupole-Orbitrap MS as described above. Raw files were processed with MaxQuant version 1.5.7.4 against a database including the *Lactobacillus* genus and common contaminants. Log_10_-transformed iBAQ intensities were compared using Mann-Whitney U test and p-values were adjusted for multiple comparisons using an FDR step-down procedure.

### FGT-METAP application

We have developed FGT-METAP (Female Genital Tract Metaproteomics), an online application for exploring and mining the FGT microbial composition and function of South African women at high risk of HIV infection. The app is available at (http://immunodb.org/FGTMetap/) and allows users to repeat the analyses presented here, as well as perform additional analyses and investigation using the metaproteomic data generated in this study.

## Abbreviations

BP: biological processes
BV: Bacterial vaginosis
CC: cellular component
FDR: false discovery rate
FGT: Female genital tract
GO: gene ontology
iBAQ: intensity-based absolute quantification
LC-MS/MS: liquid chromatography-tandem mass spectrometry
MF: molecular function
OOB: out-of-bag
PCA: principal component analysis
PSA: prostate specific antigen
RF: random forest
STIs: sexually transmitted infections
WGCNA: Weighted correlation network analysis

## Declarations

### Ethics approval and consent to participate

The University of Cape Town Faculty of Health Sciences Human Research Ethics Committee (HREC REF: 267/2013) approved this study. Women ≥18 years provided written informed consent, those <18 years provided informed assent and informed consent was obtained from parents/guardians.

### Consent for publication

Not applicable.

### Availability of data and materials

The dataset supporting the conclusions of this article is available in the FGT-METAP application, http://immunodb.org/FGTMetap/. Data are available from the corresponding author upon reasonable request. The protein accession IDs provided at http://immunodb.org/FGTMetap/ reflect UniProt IDs of each protein and additional information for each protein is provided based on UniProt and NCBI databases.

## Authors’ contributions

AA analyzed the data, developed the FGT-METAP application, and wrote the manuscript; MTM, CB, SD, and SB conducted some of the laboratory experiments, analyzed the data and contributed to manuscript preparation; MP, AI, NR, IA, NM, BC and DLT analyzed the data and contributed to manuscript preparation; LB, EW, MP, ZM, HG conducted some of the laboratory experiments and contributed to manuscript preparation; NM and JMB supervised the data analysis and contributed to manuscript preparation; AG supervised the development of FGT-METAP and hosting the application and contributed to manuscript preparation; LGB managed the clinical site for the WISH study, collected some of the clinical data and contributed to manuscript preparation; HJ supervised the data collection, analyzed the data and contributed to manuscript preparation; JSP was Principal Investigator of the WISH cohort, supervised the collection of some of the data and contributed to manuscript preparation; LM conceptualized the study, supervised the data collection, analyzed some of the data and wrote the manuscript.

## Acknowledgements

We would like to acknowledge the participants of the Women’s Initiative in Health Study.

## Supplementary table titles and legends

**Additional file 1: Table S1** Demographic and clinical characteristics of study participants stratified by female genital tract pro-inflammatory cytokine levels

**Additional file 2: Table S2** List of protein BLAST results against UniProt and NCBI nr databases (DBs). Liquid chromatography-tandem mass spectrometry was used to evaluate the metaproteome in lateral vaginal wall swabs from 113 women from Cape Town, South Africa. Primary assigned taxa, obtained based on MaxQuant version 1.5.7.4, are presented in the second column. The curated taxa (third column) were obtained after considering the top similar and frequent BLAST hits for each protein. IDs: UniProt Identity codes

**Additional file 2: Table S3** List of differentially abundant taxa distinguishing women with low and high inflammation. Liquid chromatography-tandem mass spectrometry was used to evaluate the metaproteome in lateral vaginal wall swabs from 113 women from Cape Town, South Africa. The false discovery rate (FDR) corrected p-values were obtained based on the aggregated log_2_-transformed intensity-based absolute quantification of all proteins assigned to the same taxa applying the moderated t-test (limma R package), after adjusting for confounding variables including age, contraceptives, prostate specific antigen (PSA) and sexually transmitted infections (STIs). Taxa with FDR adjusted p-values ≤0.05 and log_2_-transformed fold change ≥ 1.2 or ≤ −1.2 were considered as differentially abundant taxa.

**Additional file 2: Table S4** List of differentially abundant proteins distinguishing women with low, medium and high inflammation. Liquid chromatography-tandem mass spectrometry was used to evaluate the metaproteome in lateral vaginal wall swabs from 113 women from Cape Town, South Africa. The false discovery rate (FDR) corrected p-values were obtained based on the log_2_-transformed intensity-based absolute quantification of each protein applying the moderated t-test (limma R package), after adjusting for confounding variables including age, contraceptives, prostate specific antigen (PSA) and sexually transmitted infections (STIs). Proteins with FDR adjusted p-values ≤0.05 and log_2_-transformed fold change ≥ 1.2 or ≤ −1.2 were considered as differentially abundant proteins. IDs: UniProt Identity codes

**Additional file 2: Table S5** List of differentially abundant molecular function (MF) gene ontology terms distinguishing women with low and high inflammation. Liquid chromatography-tandem mass spectrometry was used to evaluate the metaproteome in lateral vaginal wall swabs from 113 women from Cape Town, South Africa. The false discovery rate (FDR) corrected p-values were obtained based on the aggregated log_2_-transformed intensity-based absolute quantification of all proteins assigned to the same MF term applying the moderated t-test (limma R package), after adjusting for confounding variables including age, contraceptives, prostate specific antigen (PSA) and sexually transmitted infections (STIs). Taxa with FDR adjusted p-values ≤0.05 and log_2_-transformed fold change ≥ 1.2 or ≤ −1.2 were considered as significantly abundant MFs.

**Additional file 1: Table S6** Details of MaxQuant version 1.5.7.4 parameters used for analysis of metaproteomic data obtained from lateral vaginal wall swabs from 113 women from Cape Town, South Africa.

