## Additional file 1 for "Microbial function and genital inflammation in young South African women at high risk of HIV infection"

**Table S1** Demographic and clinical characteristics of study participants stratified by female genital tract pro-inflammatory cytokine levels

| | **Low pro-inflammatory cytokines** | **Medium pro-inflammatory cytokines** | **High pro-inflammatory cytokines** |
|---|---|---|---|
| | **n/N (%)** | **n/N (%)** | **n/N (%)** |
| **Median age in years (range)** | 18 (16-22) | 19 (16-22) | 18 (16-22) |
| ***Chlamydia trachomatis*** | 4/23 (17.4) | 23/57 (40.4) | 10/32 (31.3) |
| ***Neisseria gonorrhoeae*** | 2/23 (8.7) | 4/57 (7.0) | 0/32 (0) |
| ***Trichomonas vaginalis*** | 2/23 (8.7) | 1/57 (1.8) | 3/32 (9.4) |
| ***Mycoplasma genitalium*** | 0/23 (0) | 5/57 (8.8) | 2/32 (6.3) |
| **HSV-2 (PCR positive)** | 0/23 (0) | 2/57 (3.5) | 2/32 (6.3) |
| **Yeast cells detected** | 2/23 (8.7) | 6/56 (10.7) | 5/32 (15.6) |
| **BV positive** | 3/23 (13.0) | 29/56 (51.8) | 24/32 (75.0) |
| **BV intermediate** | 2/23 (8.7) | 1/56 (1.8) | 4/32 (12.5) |
| **PSA positive** | 4/23 (17.4) | 12/57 (21.1) | 5/33 (15.2) |
| **Using DMPA** | 6/23 (26.1) | 9/57 (15.8) | 5/33 (15.2) |
| **Using Nur-Isterate** | 14/23 (60.1) | 41/57 (71.9) | 23/33 (69.7) |
| **Using OCP** | 3/23 (13.0) | 2/57 (3.5) | 1/33 (3.0) |
| **Using Implanon** | 0/23 (0) | 5/57 (8.8) | 3/33 (9.1) |
| **Using Nuvaring** | 0/23 (0) | 0/57 (0) | 1/33 (3.0) |

HSV-2: Herpes simplex virus type 2; BV: Bacterial vaginosis; PCR: Polymerase chain reaction; PSA: Prostate specific antigen; DMPA: Depomedroxyprogesterone acetate; OCP: Oral contraceptive pill

**Table S6** Details of MaxQuant version 1.5.7.4 parameters used for analysis of metaproteomic data obtained from lateral vaginal wall swabs from 113 women from Cape Town, South Africa.

| MaxQuant Parameter | Value/Description |
|---|---|
| Version | 1.5.7.4 |
| Fixed modifications | Carbamidomethyl (C) |
| Include contaminants | TRUE |
| PSM FDR | 0,01 |
| Protein FDR | 0,01 |
| Site FDR | 0,01 |
| Use Normalized Ratios For Occupancy | TRUE |
| Min. unique peptides | 0 |
| Min. razor peptides | 1 |
| Min. peptides | 1 |
| Modifications included in protein quantification | Oxidation (M);Acetyl (Protein N-term) |
| Discard unmodified counterpart peptides | TRUE |
| iBAQ | TRUE |
| iBAQ log fit | TRUE |
| Decoy mode | revert |
| Include contaminants | TRUE |
| Second peptides | TRUE |
| Stabilize large LFQ ratios | TRUE |
| Require MS/MS for LFQ comparisons | TRUE |
| Min. peptide length for unspecific search | 8 |
| Max. peptide length for unspecific search | 25 |
| Razor protein FDR | TRUE |

FDR: false discovery rate; iBAQ: intensity-based absolute quantification; LFQ: label-free quantification; min: minimum; MS/MS: tandem mass spectrometry

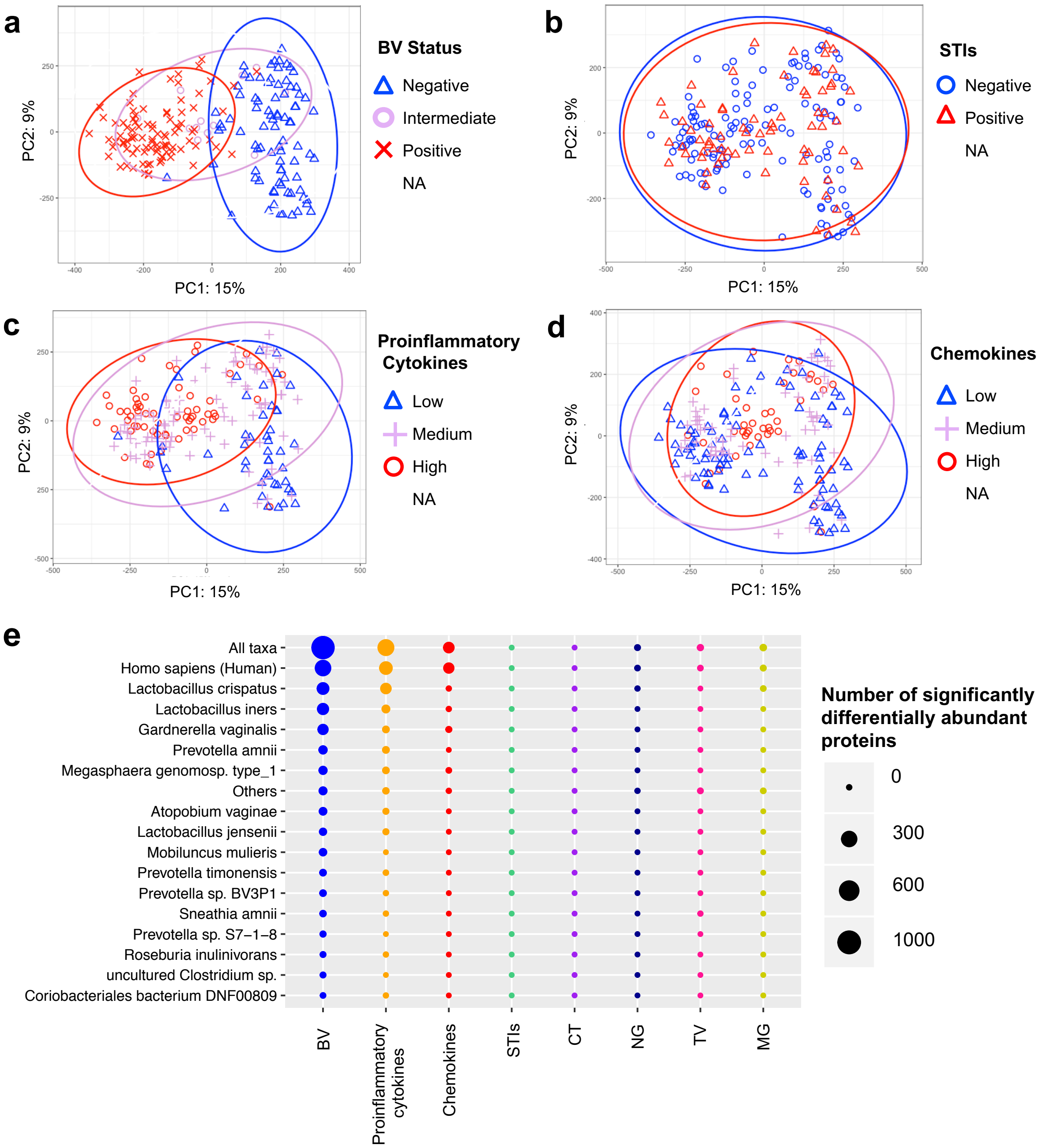


**Figure S1 Clustering of women according to overall metaproteomic profiles.** Liquid chromatography-tandem mass spectrometry was used to evaluate the metaproteome in lateral vaginal wall swabs from 113 women from Cape Town, South Africa. Principal component analysis (mixOmics R package) was used to group women based on the $log_2$-transformed intensity-based absolute quantification (iBAQ) intensities of all proteins identified. Grouping by (**a**) bacterial vaginosis (BV) status, (**b**) sexually transmitted infection (STI) status, (**c**) vaginal pro-inflammatory cytokine concentrations, and (**d**) vaginal chemokine concentrations are shown. Each point represents an individual woman. (**e**) Statistically significant differences in the relative abundance of proteins between BV positive and negative women, women with high versus low chemokine or pro-inflammatory cytokine concentrations, and STI positive versus negative women were determined using the limma package in R. Each model was adjusted for potentially confounding variables including age, contraceptives, prostate specific antigen (PSA) and co-infections. The number of statistically significant differentially abundant proteins per comparison is indicated by circle size, with the taxonomic annotation of the proteins shown on the left side of the graph. PC: Principal component. CT: *Chlamydia trachomatis*, NG: *Neisseria gonorrhoeae*, TV: *Trichomonas vaginalis* and MG: *Mycoplasma genitalium*

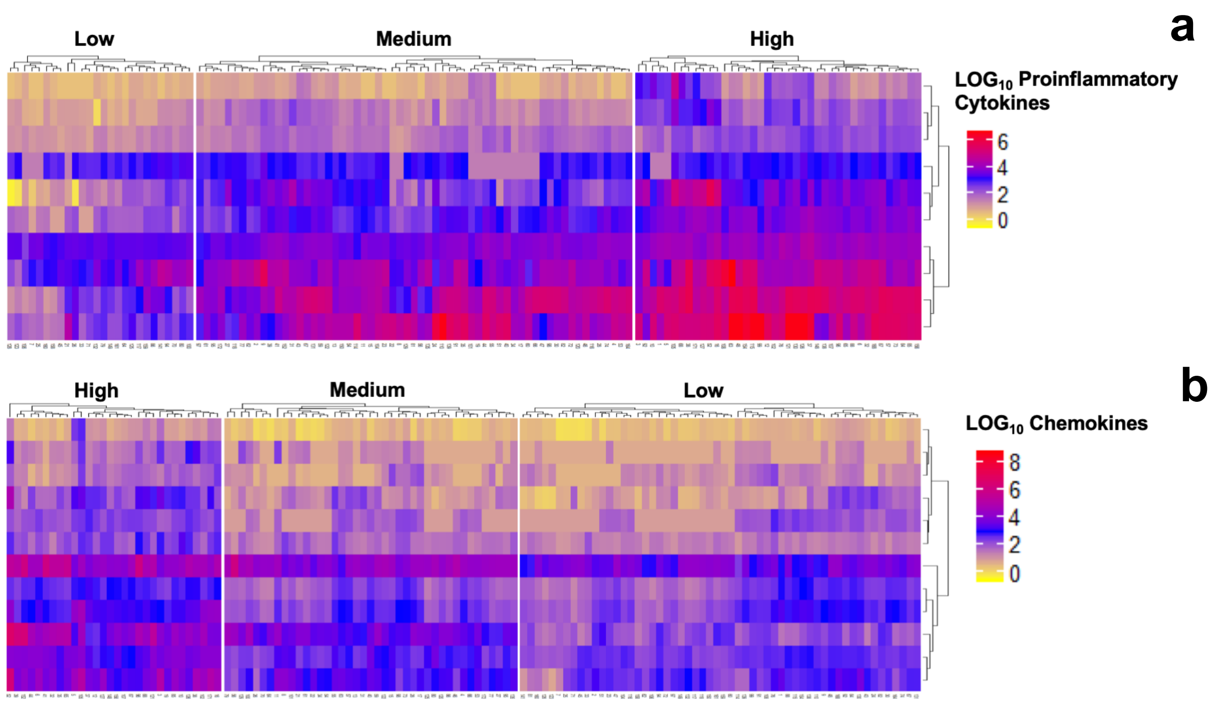


**Figure S2 Identification of women with low, medium and high levels of pro-inflammatory cytokines and chemokines.** The concentrations of 48 cytokines were measured in menstrual cup samples from all study participants (n=113). Of these cytokines, nine were classified as pro-inflammatory [interleukin (IL)-1α, IL-1β, IL-6, IL-12p40, IL-12p70, tumor necrosis factor (TNF)-α, TNF-β, TNF-related apoptosis-inducing ligand (TRAIL), interferon (IFN)-γ] and twelve were classified as chemokines [cutaneous T-cell-attracting chemokine (CTACK), eotaxin, growth regulated oncogene (GRO)-α, IL-8, IL-16, IFN-γ-induced protein (IP)-10, monocyte chemoattractant protein (MCP)-1, MCP-3, monokine induced by IFN-γ (MIG), macrophage inflammatory protein (MIP)-1α, MIP-1β, regulated on activation, normal T cell expressed and secreted (RANTES)]. Unsupervised hierarchical followed by K-means clustering were then used to identify women with low, medium and high pro-inflammatory cytokine or chemokine profiles. The heatmaps show $\log_{10}$-transformed cytokine concentrations.

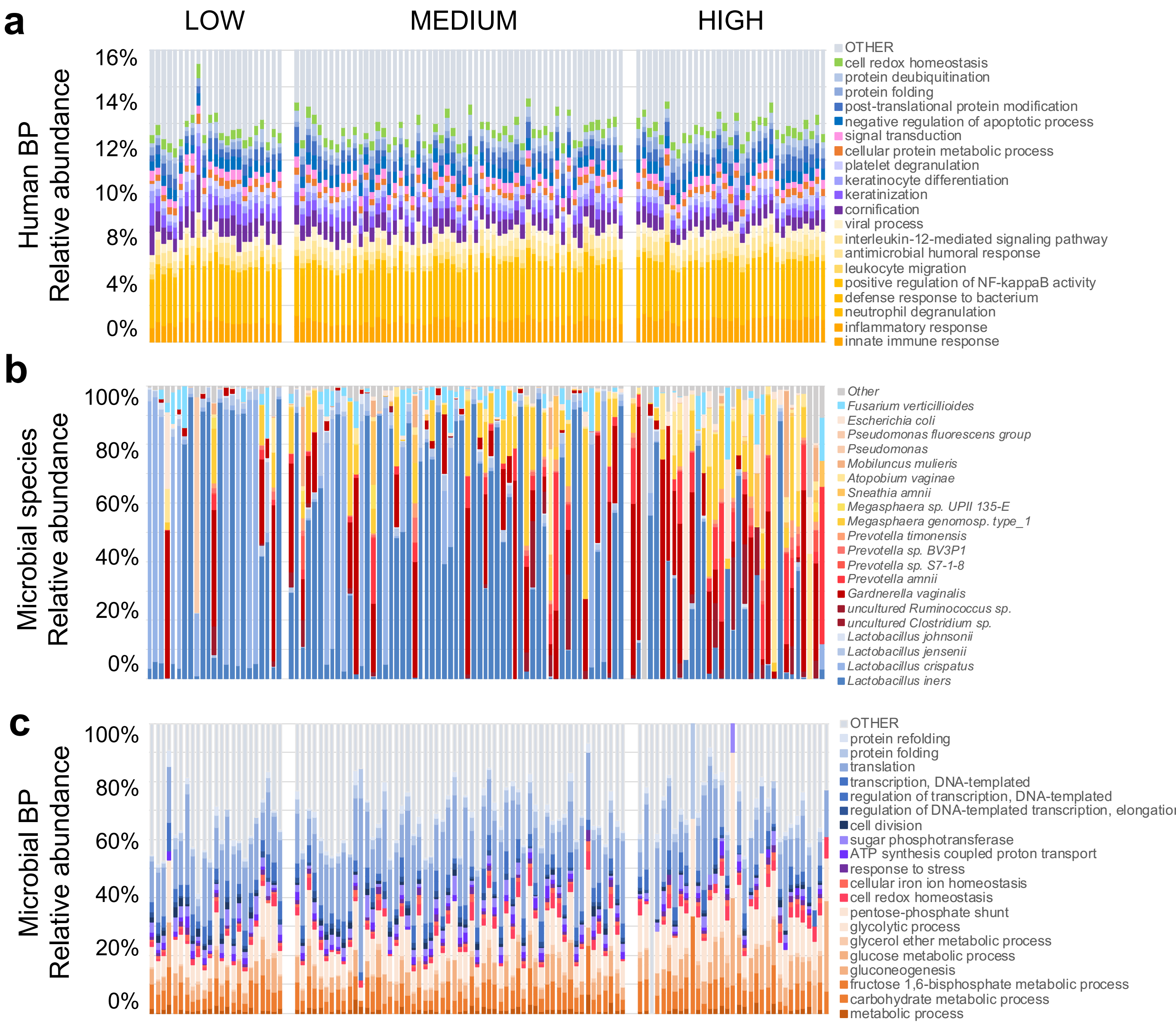


**Figure S3 Human biological processes, microbial taxa and microbial biological processes in individual women with low, medium and high inflammation.** Liquid chromatography-tandem mass spectrometry was used to evaluate the metaproteome in lateral vaginal wall swabs from 113 women from Cape Town, South Africa. Proteins were identified using MaxQuant and a custom database generated using *de novo* sequencing to filter the UniProt database. Gene ontology was determined using UniProt and the intensity-based absolute quantification (iBAQ) values of proteins with the same biological process (BP) gene ontologies were aggregated separately for human proteins and microbial proteins. The top 20 most abundant (**a**) human BPs, (**b**) microbial taxa determined using metaproteomics and (**c**) microbial BPs are shown as stacked bar graphs for individual participants. Participants are grouped according to inflammatory cytokine profiles.

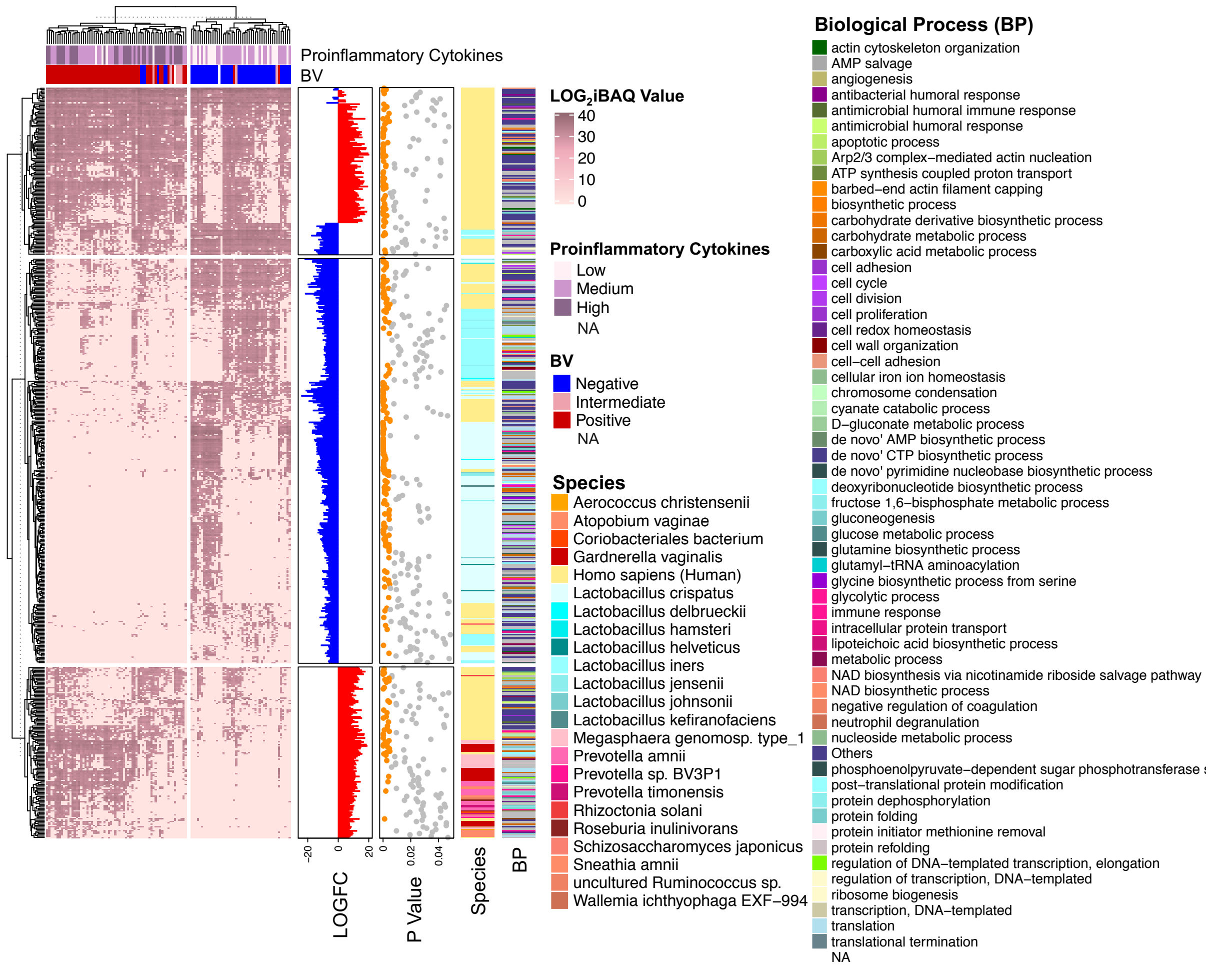


**Figure S4 Differentially abundant proteins in women with high versus low inflammatory cytokine profiles.** Unsupervised hierarchical clustering of $\log_2$-transformed intensity-based absolute quantification (iBAQ) values for significantly differentially abundant proteins followed by K-means clustering of both rows and columns was conducted using the ComplexHeatmap R package. The list of significantly differentially abundant proteins between women with low (n=23), medium (n=57) and high (n=33) inflammation was obtained using the limma R package. The p-values were calculated using the moderated t-test after adjusting for confounding variables including age, contraceptives, semen exposure and sexually transmitted infections. Proteins with false discovery rate (FDR) adjusted p-values <0.05 and $\log_2$-transformed fold change $\geq$ 1.2 or $\leq$ -1.2 were considered as significantly differentially abundant and are shown in the heatmap. The bar graph shows $\log_2$-transformed fold changes (LOGFC), with red indicating positive and blue indicating negative fold changes in women with high versus low inflammation. The species annotation and the biological process gene ontology of the proteins are shown on the right of the heatmap. Adjusted p-values are shown by dots, with orange dots indicating FDR adjusted p-values (<0.005). BV: Bacterial vaginosis. NA: Not applicable. BP: Biological Process.

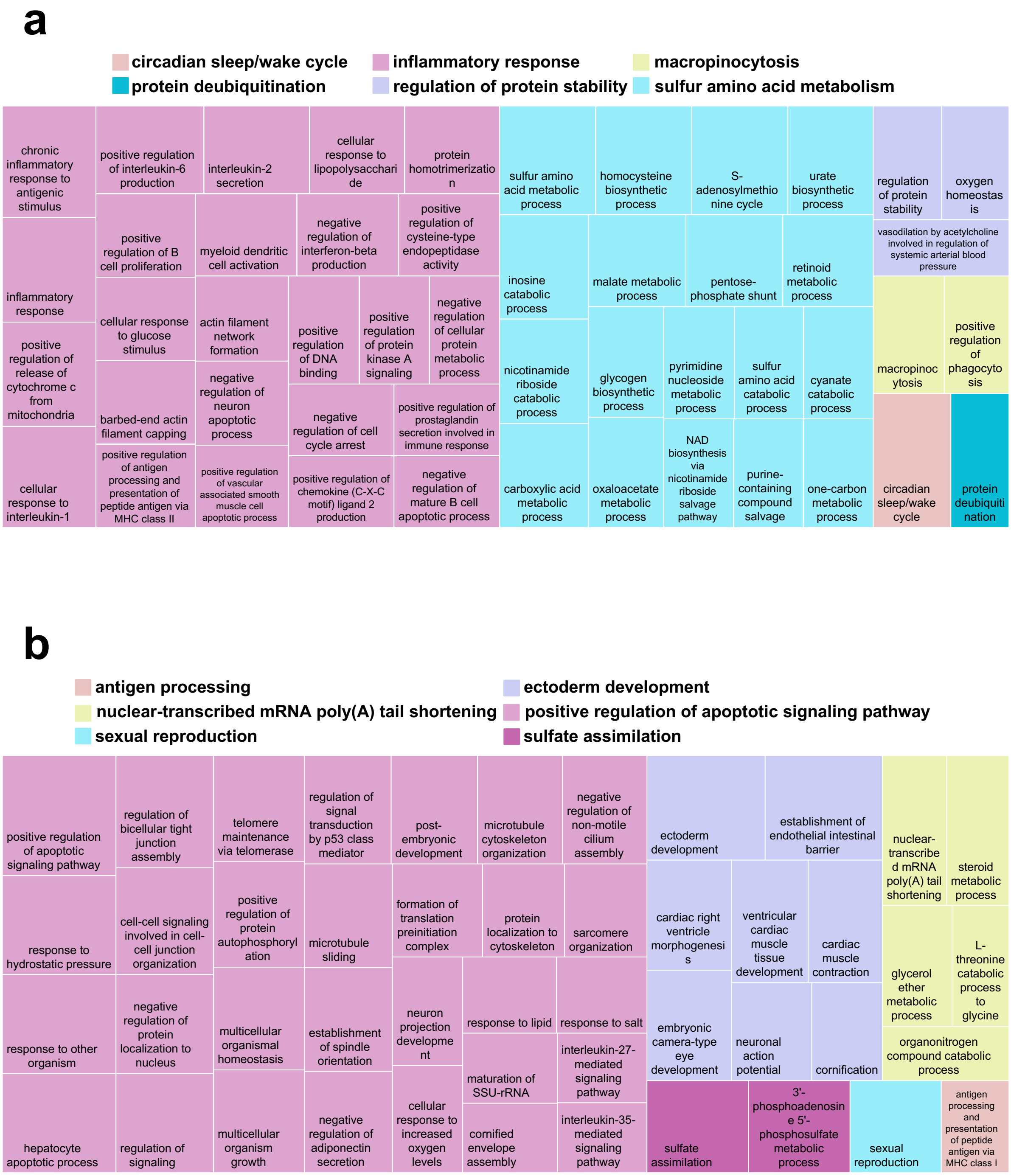


**Figure S5 Top 50 significantly differentially abundant human biological processes in women with low, and high inflammation.** Liquid chromatography-tandem mass spectrometry was used to evaluate the metaproteome in lateral vaginal wall swabs from 113 women from Cape Town, South Africa. Proteins were identified using MaxQuant and a custom database generated using *de novo* sequencing to filter the UniProt database. Gene ontology was determined using UniProt and the intensity-based absolute quantification (iBAQ) values of human proteins with the same biological process (BP) gene ontologies were aggregated. The significantly differentially abundant pathways were obtained using the moderated t-test implemented in the limma R package. False discovery rate (FDR) adjusted p-values were calculated after controlling for potentially confounding variables including age, contraceptives, semen exposure and sexually transmitted infections. Results were visualized using the REVIGO tool (http://revigo.irb.hr/). (**a**) The top 50 overrepresented human BPs in women with inflammation (FDR p-values <0.05). (**b**) The top 50 underrepresented human BPs in women with inflammation (FDR p-values <0.05).

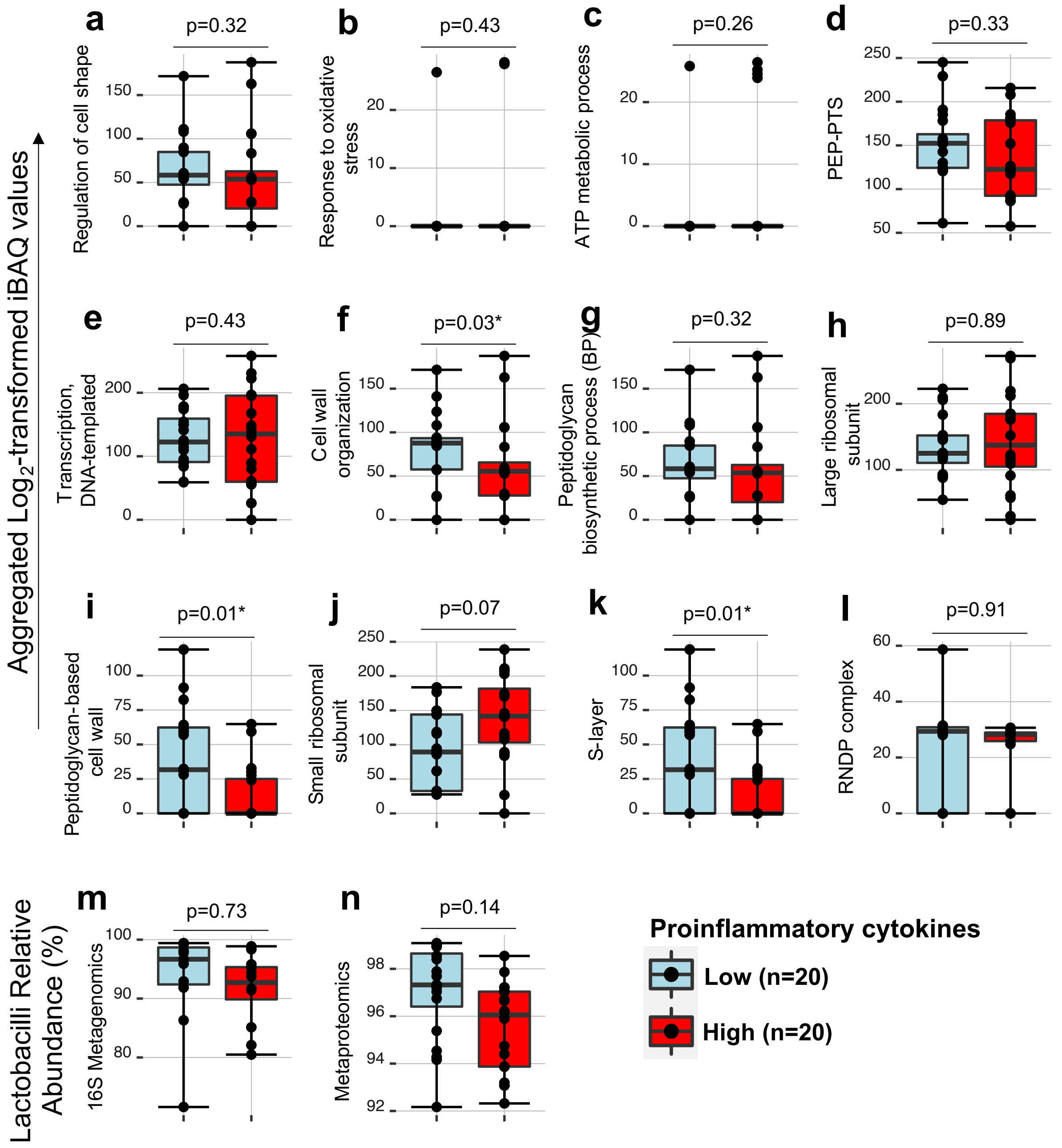


**Figure S6 Relative abundance of gene ontologies in BV negative women with *Lactobacillus* dominant communities.** To further investigate whether the associations between FGT inflammation and *Lactobacillus* biological processes (BPs) and cellular components (CCs) were independent of *Lactobacillus* relative abundance, we identified women with *Lactobacillus* dominant communities (n=40). To do this, we excluded women who were BV positive or BV intermediate, included women with *Lactobacillus* relative abundance (by metaproteomics) of >90% and lastly removed women with less than 65% *Lactobacillus* relative abundance (based on 16S metagenomics data if available). Women were then grouped as having low (n=20) versus relatively high (n=20) inflammatory cytokines by grouping all of nine proinflammatory cytokines [interleukin (IL)-1α, IL-1β, IL-6, IL-12p40, IL-12p70, tumor necrosis factor (TNF)-α, TNF-β, TNF-related apoptosis-inducing ligand (TRAIL), interferon (IFN)-γ] onto one principal component and generating component estimates for each woman. Women with component estimates above the median were considered to have relatively high proinflammatory cytokines, while those below the median were considered to have low proinflammatory cytokines. The aggregated iBAQ values for the top 12 BPs and CCs distinguishing women with low versus high inflammatory cytokines in the full cohort were then compared in this sub-cohort of women with *Lactobacillus* dominant communities. The p-values were obtained using the moderated t-test after adjusting for confounding variables including age, contraceptives, prostate specific antigen (PSA), and sexually transmitted infections (STIs).

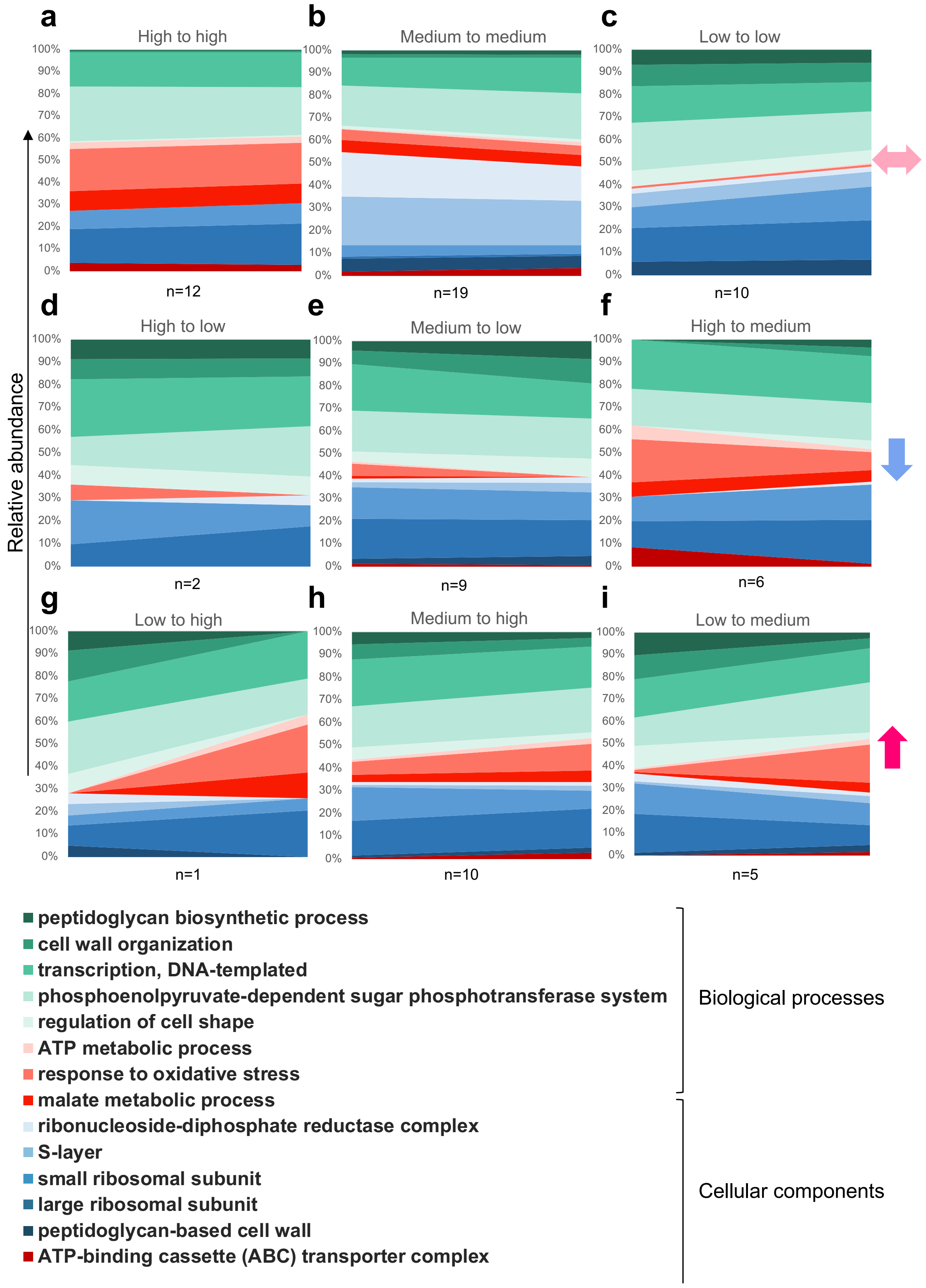


**Figure S7 Longitudinal changes in FGT microbial biological process and cellular component gene ontologies.** Liquid chromatography-tandem mass spectrometry was used to evaluate the metaproteome in lateral vaginal wall swabs from 74 women from Cape Town, South Africa in two visits 9 weeks apart. Inflammation groups at both visits were defined based on hierarchical followed by K-means clustering of nine pro-inflammatory cytokine concentrations [interleukin (IL)-1α, IL-1β, IL-6, IL-12p40, IL-12p70, tumor necrosis factor (TNF)-α, TNF-β, TNF-related apoptosis-inducing ligand (TRAIL), interferon (IFN)-γ]. Top 14 biological process and cellular component gene ontology terms distinguishing women with and without inflammation at both visits were determined by moderated t-test (limma R package) and random forest algorithm (randomForest R package). Changes in relative abundance of these biological processes and cellular components are shown as area plots.
